# VIOLIN: A modular framework for scalable reconciliation of heterogeneous interaction graphs

**DOI:** 10.1101/2024.07.21.604448

**Authors:** Haomiao Luo, Casey Hansen, Niloofar Arazkhani, Cheryl A. Telmer, Difei Tang, Gaoxiang Zhou, Peter Spirtes, Natasa Miskov-Zivanov

**Affiliations:** University of Pittsburgh, Pittsburgh PA; Carnegie Mellon University, Pittsburgh PA

## Abstract

Automated extraction of molecular interactions from scientific literature has outpaced the development of systematic methods for integrating this information with curated models and knowledge graphs. Here we present VIOLIN (Versatile Interaction Organizing to Leverage Information in Networks), a configurable, attribute-aware reconciliation framework that formally compares newly extracted interaction lists against structured baseline graphs. VIOLIN classifies each interaction as a corroboration, contradiction, flagged case, or extension, and supports configurable attribute inclusion strategies and mismatch semantics to adjust reconciliation strictness. We evaluate VIOLIN using interaction lists generated by two traditional NLP systems (REACH, INDRA) and two large language models (GPT-4.1, Llama 3) across multiple literature corpora and structurally distinct baseline graphs. Across all conditions, reconciliation outcomes were stable and interpretable, with extensions dominating and corroboration–contradiction balance reflecting intrinsic structural relationships between baseline graphs and extracted evidence. Sensitivity analyses demonstrate that attribute inclusion and classification scheme selection shift category boundaries predictably. Benchmark evaluations confirm high algorithmic correctness and alignment with expert curation. VIOLIN is publicly available as a Python package and through web-based interface (https://nmzlab.github.io/Tools-UI).

## 1 Introduction

Mechanistic knowledge in biology is often represented as structured interaction networks, where nodes correspond to molecular entities and edges encode directed, signed regulatory relationships^1–11^. Such network representations form the basis of executable models used to simulate cellular behavior, reason about perturbations, and generate experimentally testable hypotheses^12–22^. At the same time, advances in natural language processing (NLP) and large language models (LLMs) have dramatically accelerated the extraction of molecular interactions from the scientific literature^23–28^. As a result, new interactions can now be generated at a scale that far exceeds the capacity of manual model creation.

Despite these advances in extraction, systematic integration of newly retrieved interactions into existing knowledge graphs and mechanistic models remains a major bottleneck. Extracted interactions differ in structure, granularity, and attribute completeness depending on the underlying extraction system and paper corpus. Moreover, mechanistic models are often context-specific, attribute-rich, and curated with particular modeling objectives in mind. As a result, simply appending newly extracted interactions to an existing network is insufficient and may introduce redundancy, contradiction, or biologically implausible structure. What is needed is a formal reconciliation framework that can compare newly extracted knowledge against a baseline graph and determine, in a structured and reproducible manner, whether each interaction corroborates, contradicts, extends, or ambiguously relates to existing model structure.

Current approaches to updating mechanistic models^1,2,9,10^ typically rely on manual curation or ad hoc filtering rules^29,30^. While database-backed validation tools can remove unsupported interactions, they do not formally distinguish structural corroboration from contextual disagreement, nor do they expose alternative semantic interpretations of mismatches^30–39^. Conversely, literature extraction tools can be used for generating candidate interactions but do not provide mechanisms for evaluating those interactions relative to a target knowledge graph or executable model^23–25,28^. The absence of a configurable, attribute-aware reconciliation engine limits the scalability and reproducibility of literature-to-model workflows.

Here we present VIOLIN (Versatile Interaction Organizing to Leverage Information in Networks), a graph reconciliation framework designed to formally compare a new interaction list against a baseline interaction network. VIOLIN defines explicit element and interaction match conditions, incorporates path-based structural reasoning, and classifies each candidate interaction into one of four primary categories: corroboration, contradiction, extension, or flagged. Importantly, VIOLIN treats reconciliation as a configurable decision process. It supports alternative classification schemes that alter how connection and path-level mismatches are interpreted, as well as attribute inclusion strategies that control which contextual attributes influence classification outcomes. This parameterization enables users to adjust semantic strictness according to modeling objectives without modifying the underlying graph structure.

We evaluate VIOLIN using heterogeneous interaction lists generated by both traditional NLP-based systems and LLMs across multiple literature corpora. By reconciling these interaction sets against structurally distinct biological knowledge graphs, we examine how extraction variability, attribute completeness, and corpus design influence reconciliation outcomes. This experimental design treats extraction systems as structured perturbations to the reconciliation process, enabling systematic evaluation of robustness across diverse input conditions.

Across all corpora and extraction strategies, VIOLIN produces stable and interpretable classification distributions, with extensions representing structural novelty, corroborations supporting existing model structure, and contradictions highlighting potential conflicts or new regulatory hypotheses. Sensitivity analyses demonstrate how attribute inclusion and classification scheme selection shift category boundaries in predictable ways, revealing the importance of explicit semantic configuration. Integration with external filtering tools further illustrates how database-supported validation interacts with graph-based reconciliation.

## 2 Results

### 2.1 Framework for structured reconciliation of extracted knowledge with baseline graphs

We developed VIOLIN as a configurable reconciliation framework that aligns newly extracted interaction lists with structured baseline knowledge graphs (Figure 1). The framework accepts two inputs: (i) a baseline knowledge graph representing curated interaction knowledge and (ii) a newly extracted interaction list derived from literature, databases, or manual curation. The output is a classified interaction list in which each newly extracted interaction is evaluated relative to the baseline and assigned to one of four categories: corroboration, contradiction, flagged, or extension.

**Figure 1.**
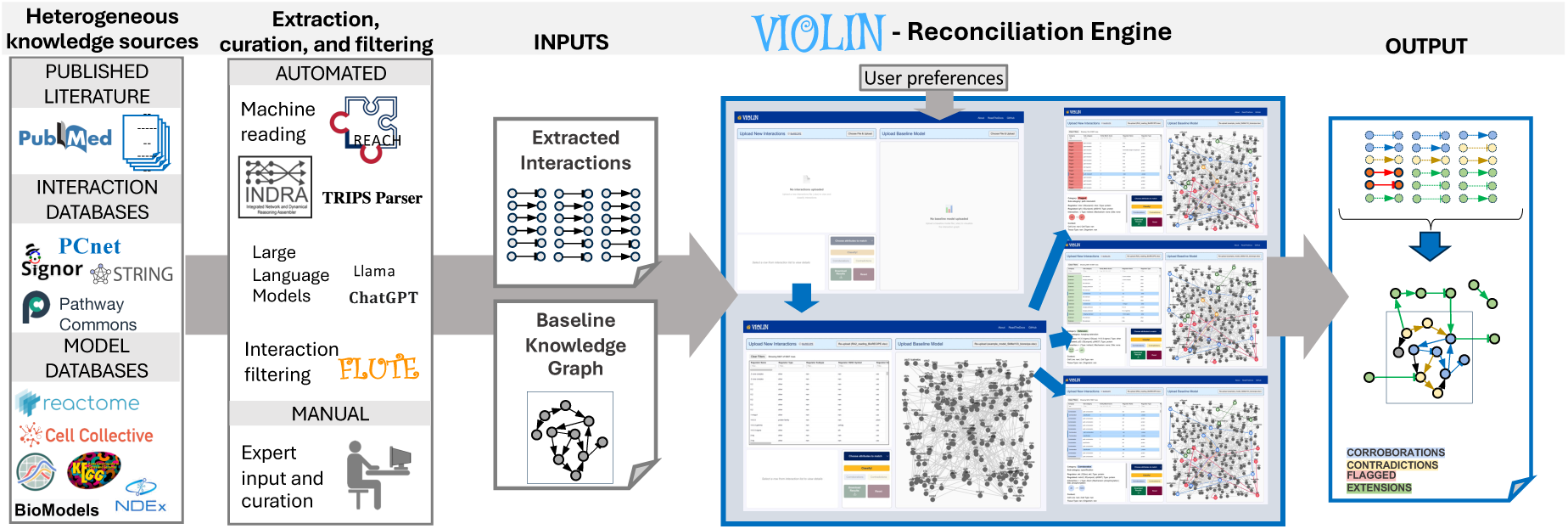
Overview of the workflow for VIOLIN with its input sources and types, and its output. Heterogeneous knowledge sources, including literature-derived extractions and curated knowledge graphs and mechanistic models, can be used as inputs to the VIOLIN reconciliation engine. The reconciliation engine performs attribute-aware graph comparison using configurable decision schemes and path-level reasoning to classify interactions as corroborations, contradictions, flagged cases, or extensions. User-defined attribute inclusion strategies and filtering modules allow controlled sensitivity analysis and workflow customization. All listed databases and tools are compatible with VIOLIN either directly or through format converters.

New interaction lists may originate from heterogeneous sources, including rule-based NLP systems^40^, structured databases^7,41–43^, LLMs^44,45^, or expert-curated corpora. Baseline graphs may be assembled from curated pathway repositories^1,2,46^, executable mechanistic models^9,47,48^, or knowledge graph and mechanistic model databases^8,49^. By separating extraction from reconciliation, VIOLIN functions as a modular infrastructure component capable of integrating diverse upstream knowledge sources into structured evaluation workflows.

To ensure interoperability and reproducibility, VIOLIN operates on interaction graphs encoded in the BioRECIPE format^48,50,51^, a standardized representation for node identity, edge directionality and sign, and contextual attributes (Supplement Section 1.1). This abstraction enables consistent attribute-aware comparison across heterogeneous inputs while maintaining compatibility with model development and analysis platforms^32,34,36–38,52–57^. Since the framework relies on directed interaction graphs, it can be generalized to diverse biological networks.

The classified output supports multiple downstream goals. Corroborations provide automated verification of baseline interactions, contradictions highlight potential conflicts or knowledge gaps, flagged cases identify ambiguous comparisons requiring manual inspection, and extensions represent candidate interactions for model expansion. In this way, VIOLIN formalizes the transfer of knowledge from additional or new sources into structured graph representation for further analysis and predictions.

### 2.2 Formalization of systematic attribute-aware graph comparison

VIOLIN evaluates new interactions by systematically comparing their attributes to those of baseline graph edges using a set of formal match definitions (Methods, Supplement Section 1.2). The reconciliation algorithm operates at the level of nodes, directed signed edges, and multi-edge paths (Figure 2A). Element matching requires agreement in unique element identifiers and type, while interaction matching requires consistency in sign and source–target mapping. Additional attributes, including compartment, mechanism and contextual metadata, are incorporated through configurable decision rules.

**Figure 2.**
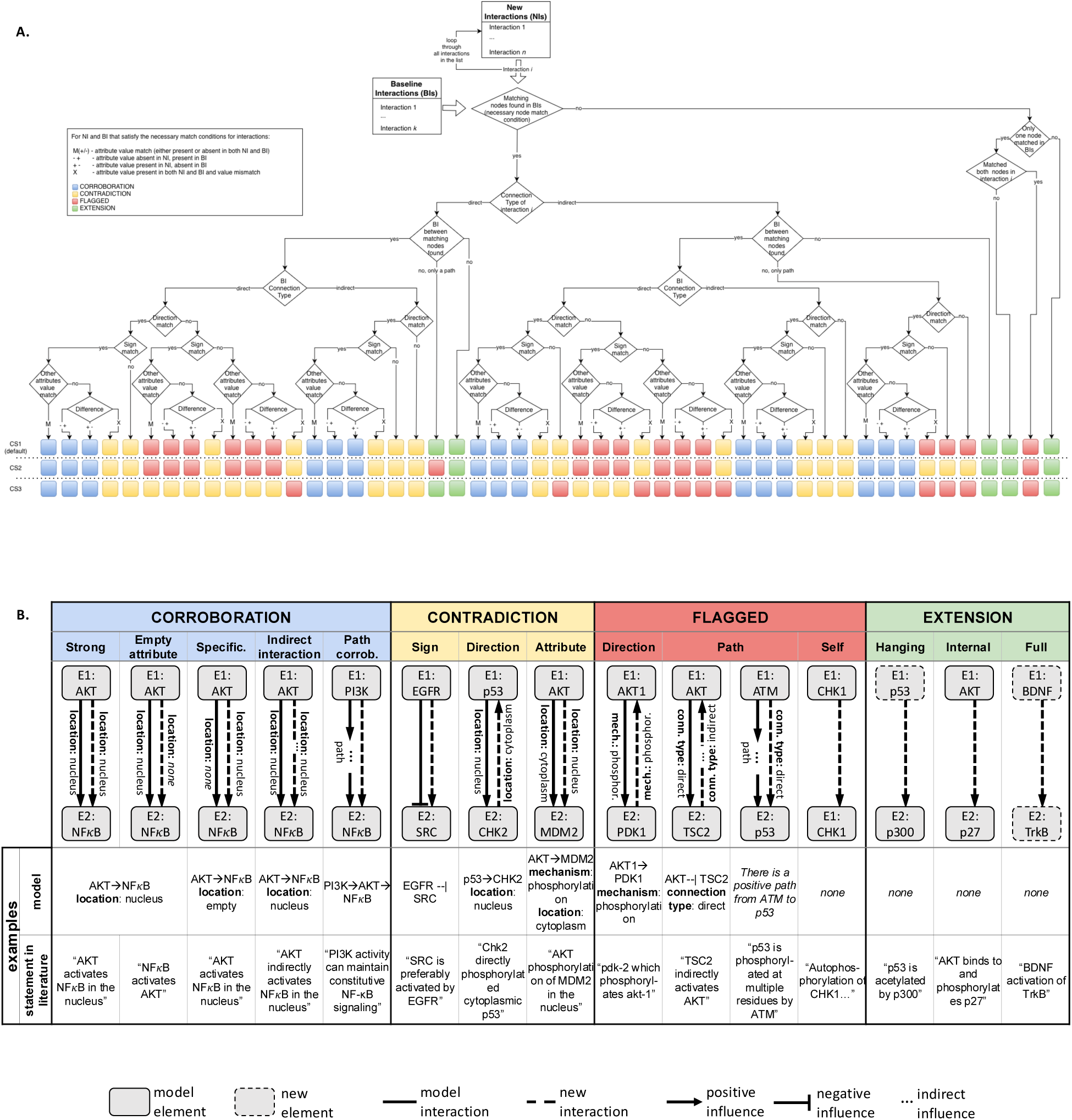
VIOLIN’s classification method, categories, schemes, and examples. **A.** VIOLIN’s decision algorithm with three classification schemes, CS1 (default), CS2 and CS3. **B.** Examples and illustrations of classification categories and sub-categories.

Attribute mismatches are detected when corresponding attributes differ between matched interactions, excluding cases where attribute values are absent. Provenance attributes (e.g., source document identifiers) are not included in classification logic to ensure that reconciliation decisions reflect structural and mechanistic consistency rather than citation overlap.

The framework supports multiple classification schemes (Methods) that enable exploration of how alternative modeling assumptions influence reconciliation outcomes. Schemes CS1-CS3 (Figure 2A) allow users to adjust how connection mismatches and path-level inconsistencies are interpreted. For example, direction mismatches may be interpreted as contradictions under one scheme and flagged under another, providing flexibility for distinct modeling objectives. The sensitivity of output to different classification decisions is detailed in Sections 2.8 and 2.9.

Within each of the four primary categories, VIOLIN further distinguishes several subcategories (Figure 2B). Corroborations distinguish between exact attribute matches, missing attributes, indirect interaction matches, path-level confirmations, and specifications. Contradictions encompass sign, direction, or attribute conflicts. Extensions are classified based on their structural relationship to the baseline graph, and can be disconnected, hanging, or internal to the baseline graph. Flagged interactions capture ambiguous comparisons that do not satisfy criteria for the other categories. Category and subcategory definitions are provided in Supplement Sections 1.3-1.6. By implementing reconciliation as attribute-aware graph comparison with configurable semantics, VIOLIN provides a transparent and reproducible mechanism for integrating heterogeneous interaction information.

### 2.3 Impact of extraction heterogeneity on interaction list structure

To evaluate VIOLIN’s reconciliation behavior when different extraction methods are used, we collected several paper corpora and generated interaction lists (Methods, Supplement Section 2) using traditional NLP-based systems REACH^23^ and INDRA^28^, and LLMs GPT-4.1 and Llama 3. These systems differ substantially in extraction throughput, attribute completeness, and contextual representation (Figure 3, Supplement Section 3), and therefore, provide a comprehensive test of VIOLIN’s robustness across heterogeneous inputs.

**Figure 3.**
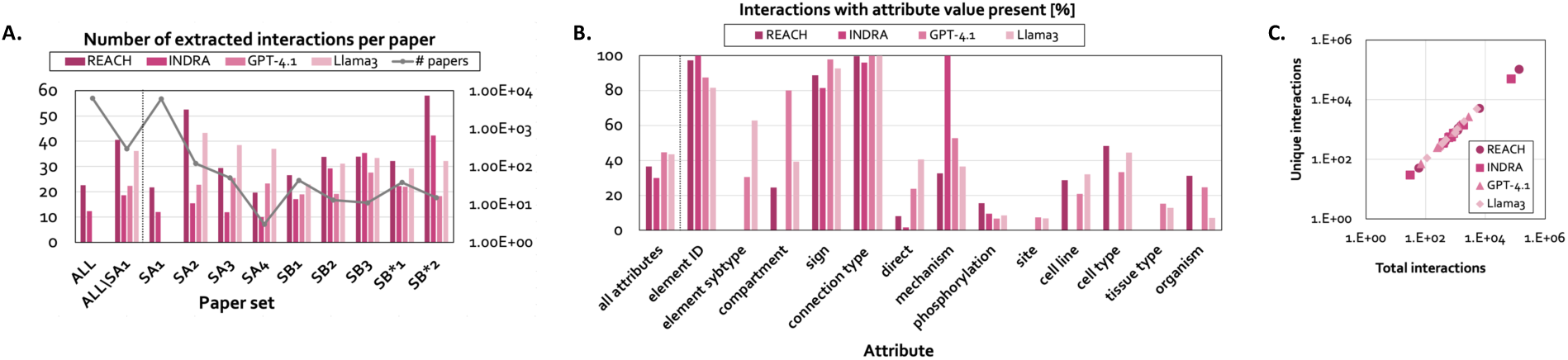
Overview of the information obtained by the four used reader tools for all corpora. **A.** The number of extracted interactions per paper across all paper corpora for each reader (ALL), across all corpora except S_A1_ since this corpus was not processed by GPT-4.1 and Llama 3 due to significant time and resource requirement (ALL\SA1), and the number of interactions per paper for each reader and each individual corpus. The line (#papers) indicates the number of papers in each corpus. **B.** The % of interactions with non-empty value for each attribute, across all interactions lists (R_A1-4_, R_B1-3_, and R_B*1-2_) for each reader and the overall presence of attributes for each reader (all attributes). **C.** The number of unique interactions vs. total interactions extracted by each reader for each corpus.

Extraction throughput varied across both readers and paper corpora (Figure 3A). Rule-based systems exhibited divergent behaviors: overall, REACH produced the highest interaction counts per document, whereas INDRA was most conservative. LLM-based systems demonstrated comparatively stable extraction densities across corpora, mainly reflecting differences in generative architecture and contextual inference. Runtime per document also differed across readers, with database-integrated NLP systems executing more rapidly than LLM-based systems (Figure S3B).

Beyond extraction density, readers differed in attribute completeness (Figure 3B). REACH shows stronger performance in finding element IDs, cell types, and organism, while INDRA performs best on extracting interaction mechanism. LLM-based systems perform strongly across most attributes and often surpass the traditional NLP-based tools. These attribute distribution differences represent distinct structural biases in the resulting interaction lists.

Importantly, we observed a strong correlation between total extracted interactions and the number of unique interactions across readers, indicating that increased extraction density generally increases structural graph diversity and is not merely a reflection of repeated interaction mentions in literature (Figure 3C).

The heterogeneous interaction corpora and extraction systems enabled analysis of how attribute completeness, extraction density, and contextual variability propagate through VIOLIN’s reconciliation framework and shape classification outcomes.

### 2.4 Global reconciliation patterns across heterogeneous inputs

We next examined how interactions extracted by heterogeneous readers were classified relative to the baseline graphs (Figure 4). To evaluate reconciliation behavior across both structurally distinct network contexts and heterogeneous inputs, we selected several curated baseline graphs that differ in size, topology, and attribute composition (Methods, Supplement Section 4). Across all extraction systems and corpora, extensions constituted the majority of classified interactions. Corroborations, contradictions, and flagged cases comprised smaller but consistently present fractions of the interaction lists, with corroborations typically occurring slightly more frequently than contradictions or flagged interactions.

**Figure 4.**
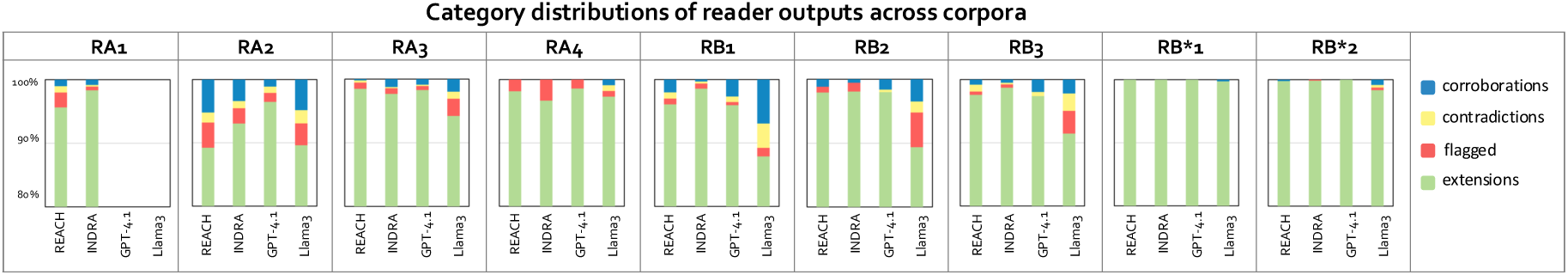
Global classification distributions. Normalized distributions of classification categories (Corroborations, Contradictions, Flagged, and Extensions) for all nine interaction lists (R_A1-4_, R_B1-3_, and R_B*1-2_), and all four readers (REACH, INDRA, GPT-4.1 and Llama 3).

Although high-level category distributions were broadly similar, some differences emerged at the level of baseline and corpus-specific graph topologies. Interaction lists related to densely connected regions of the baseline graph exhibited greater diversity across classification categories, whereas lists corresponding to sparsely connected regions and those not relevant to baseline model were dominated by extensions. This pattern indicates that reconciliation outcomes are influenced by baseline graph connectivity and corpus heterogeneity rather than corpus size alone. Differences across extraction systems were also evident for each interaction list (Figure 4). Notably, interaction graphs generated by Llama 3 consistently exhibit the most significant diversity across reconciliation categories, reflecting increased attribute richness and contextual variability in its output. These differences indicate that extraction outputs can be treated as structured perturbations of the input graph space when evaluating reconciliation behavior.

### 2.5 Structural interpretation of reconciliation subtypes

To better understand reconciliation dynamics, we analyzed the distribution of subcategories within the four primary classification categories (Figure S5).

For corroborations, path-level confirmations were the dominant subtype across readers and corpora. These interactions did not correspond to direct edge matches but aligned with multi-edge paths in the baseline graph, indicating that literature-derived evidence frequently reinforces indirect regulatory relationships. Distributions of other corroboration subtypes varied across extraction systems. Traditional NLP-based readers more frequently produced indirect corroborations, consistent with their conservative assignment of direct interaction types (Figure S6). LLM-derived interaction lists showed a more balanced distribution of corroboration subtypes, including strong and specification corroborations, which reflects greater attribute completeness and contextual specificity. In particular, LLMs more frequently populated interaction attributes beyond those used in minimal match requirements, enabling more precise alignment classifications.

Contradictions were primarily driven by sign mismatches for traditional NLP systems, whereas LLM-derived inputs exhibited a more balanced distribution between sign and direction mismatches. Attribute-level conflicts were comparatively rare across all readers. This distribution indicates that most disagreements between extracted and baseline graphs arise from differences in regulatory polarity or edge orientation rather than contextual metadata discrepancies.

Flagged interactions were predominantly associated with path mismatches, in which extracted interactions corresponded to baseline paths but exhibited one or more attribute inconsistencies. These ambiguous alignments highlight structural regions where additional evidence or refined modeling assumptions may be required. Additionally, these interactions may also indicate that new feed-forward or feedback loops could be formed in the baseline graph.

Overall, these subtype patterns further support the observation that reconciliation behavior is affected not only by reader characteristics but also by graph topology and attribute presence in the baseline graph.

### 2.6 Balance between corroboration and contradiction across baseline coverage

We next examined the relative balance between corroborations and contradictions across all corpora and extraction systems (Figure 5, left). Corroborations generally exceeded contradictions in the extracted interaction lists, consistent with the expectation that curated baseline graphs are largely supported by the literature. However, this dominance was modest, and contradictions remained consistently present across datasets.

**Figure 5.**
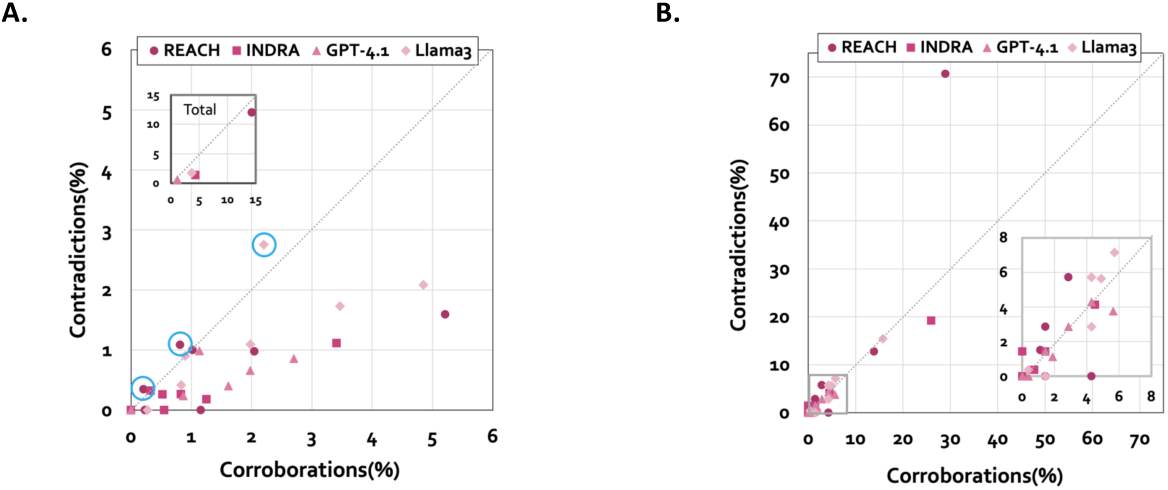
Comparison of the presence of corroborations and contradictions in interaction lists. Correlations between corroborations and contradictions in each reader output, for each corpus, presented as: **A.** Percent of the total number of interactions in the interaction list (circled markers are the only three with contradictions higher than corroborations; Inset: Overall percent across all interaction lists for each reader. **B.** Percent of the model interactions that are corroborated or contradicted; Inset: Zoomed in part of the plot for interval 0-8%.

When assessed in terms of baseline graph coverage, the relationship between corroboration and contradiction became more balanced (Figure 5, right). No extraction system achieved complete verification of the baseline graphs, even when large corpora were used. This observation suggests that baseline incompleteness, corpus scope, and extraction variability jointly limit full structural alignment.

Reader-specific trends emerged in baseline coverage, reflecting variation in extraction density and attribute completeness across systems. Llama 3 produced the largest numbers of both corroborations and contradictions and achieved the highest baseline coverage, followed by REACH. GPT-4.1 and INDRA yielded similar overall numbers of corroborations and contradictions; however, GPT-4.1 was the only reader that consistently produced a higher proportion of corroborations relative to contradictions. Across individual corpora, LLM-based readers generally provided slightly greater coverage of both corroborating and contradicting interactions than traditional NLP readers. These findings indicate that extraction density and attribute richness influence baseline verification coverage.

Importantly, contradictions identified through reconciliation may reflect either extraction artifacts or genuinely novel or context-specific evidence. The consistent presence of both corroborations and contradictions across heterogeneous corpora supports the interpretation that reconciliation captures an inherent tension between curated knowledge and evolving literature. These results demonstrate that VIOLIN enables systematic quantification of model–literature alignment, revealing both structural support and potential conflict regions within baseline graphs.

### 2.7 Structural expansion potential of extracted graphs

Across all extraction systems and corpora, extensions represented the dominant classification outcome, indicating that literature-derived interaction graphs contain substantial information not currently represented in the baseline graphs (Figure 4). This dominance persisted regardless of extraction architecture or corpus composition, suggesting that curated baseline models remain incomplete relative to literature.

Extension subtypes (Figure S5) provide further insight into graph expansion potential. Full extensions, new interactions disconnected from the baseline graph, constituted the largest extension subtype across most corpora. These interactions represent candidate knowledge that may either reflect model incompleteness or arise from grounding inconsistencies in upstream extraction. Hanging extensions, new interactions that share only one node with the baseline graph, were the second most common subtype and represent structurally adjacent opportunities for incremental graph augmentation^38^. Internal extensions, which connect existing baseline nodes with a new edge, occurred less frequently but may directly suggest new regulatory pathways within the existing graph topology.

When interaction lists were derived from intentionally weakly related or irrelevant corpora (S_B*1_ and S_B*2_), extensions were overwhelmingly classified as full extensions and rarely connected structurally to the baseline graph. This pattern confirms that the reconciliation framework appropriately distinguishes structurally unrelated input, demonstrating robustness to irrelevant literature. Conversely, when corpora were topically aligned with the baseline graph (S_A1-4_ for model A, S_B1-3_ for model B), the proportion of hanging and internal extensions increased, indicating targeted graph expansion opportunities.

Some full extensions were attributable to upstream entity grounding discrepancies, wherein equivalent biological entities were represented with different identifiers. These cases highlight the impact that varying extraction approaches can have on reconciliation classification. Nevertheless, the overall distribution of extension subtypes can assist in quantifying incompleteness of the baseline graph and systematically identifying regions for its expansion.

### 2.8 Sensitivity of classification outcomes to attribute inclusion

Although element identity and interaction sign constitute the necessary conditions for structural matching, other attributes such as compartment, mechanism, and contextual metadata can also be included in classification decisions. Since the presence of these attributes varies across reader outputs and corpora (Figure 3B, Figure S6), we systematically varied non-essential attribute inclusion using strategies CA1–CA4 (Methods) to evaluate their effect on reconciliation outcomes.

Across all strategies and extraction systems, extensions remained dominant, with more than 90% of interactions consistently classified as extensions (Figure 6A, row). Nevertheless, we observed some reclassifications when mechanism and contextual attributes were included. Mechanism information alone had relatively limited influence, causing only small shifts from corroborations to contradictions (≈1–2%). In contrast, contextual attributes, particularly cell line, produced substantially larger changes. When cell line information was incorporated (CA2→CA3), the number of corroborations decreased markedly, while flagged category substantially increased. Most of these flagged interactions correspond to path mismatches caused by contextual inconsistencies between literature-derived interactions and the baseline models.

**Figure 6.**
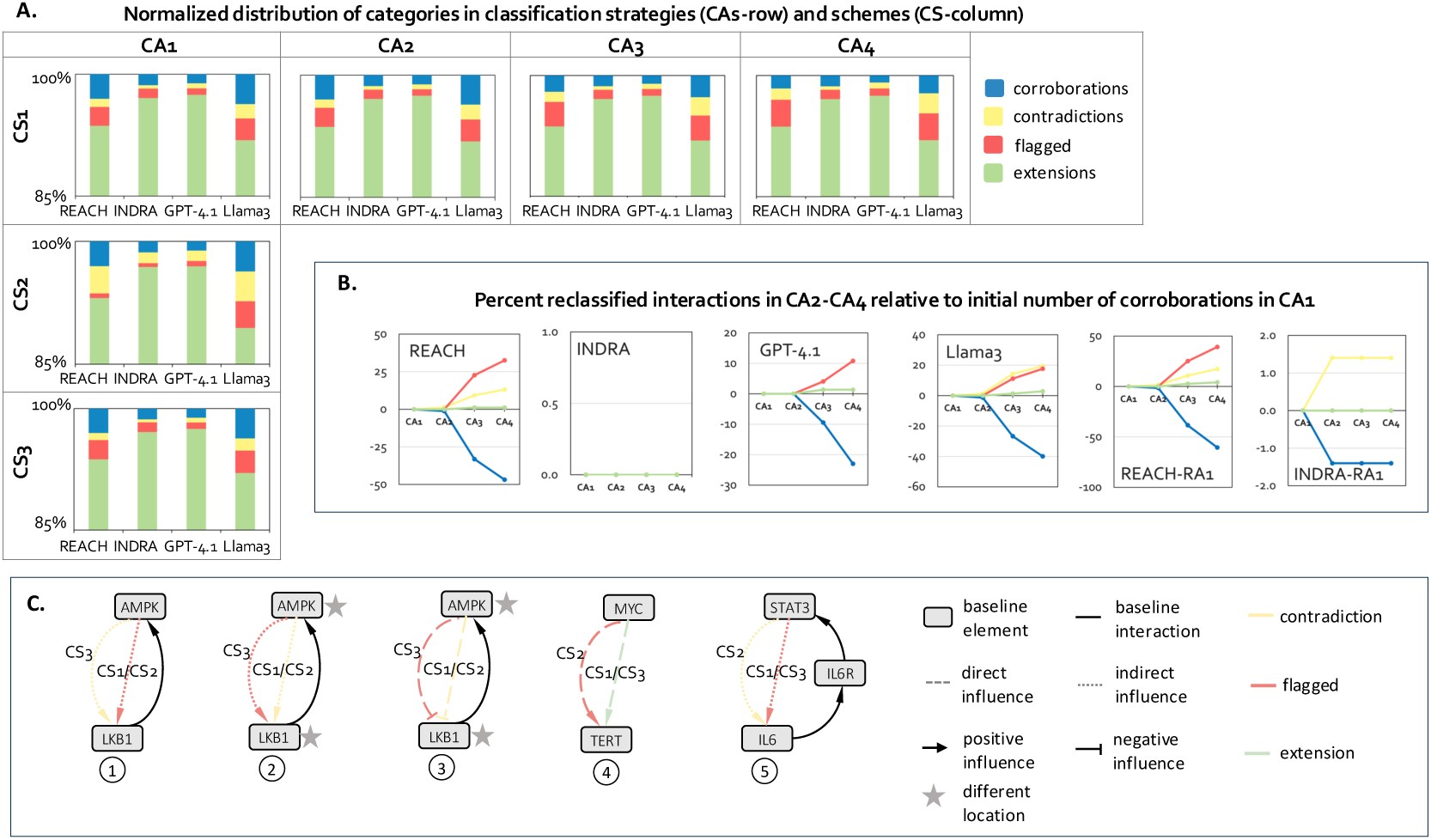
Sensitivity to attribute inclusion and to alternative mismatch semantics. **A.** (row) Sensitivity to attribute choice strategies (CA1, CA2, CA3, CA4) under classification scheme CS1; (column) sensitivity to alternative mismatch semantics (CS1, CS2, CS3) under CA1 strategy. All numbers are summed across all corpora except R_A1_ (since R_A1_ was not extracted by GPT-4.1 and Llama3). **B.** Percent change in all categories for CA2, CA3, CA4, computed relative to the total number of corroborations in CA1, for each reader across all corpora except R_A1_, and in R_A1_ for REACH and INDRA only. Corroborations get reclassified into contradictions, flagged, and extensions in CA2, CA3, and CA4. **C.** Five examples of interactions that are classified differently with CS1, CS2, and CS3 (descriptions below). (1) : New interaction indicates that AMPK has an indirect positive influence on LKB1, while in the curated baseline graph, LKB1 has a direct positive influence on AMPK – the new and baseline interaction have same sign and opposite direction. If the user prefers to consider this as a potential new interaction and further manually examine, CS1 or CS2 can be used to classify this interaction as flagged; if the opposite direction should be strictly taken as contradiction only, then CS3 can be used. (2) : New interaction indicates that AMPK has an indirect positive influence on LKB1, while in the curated baseline graph, LKB1 directly regulates AMPK to enhance its expression. Additionally, the location of AMPK and LKB1 are different between baseline graph and new interaction. The new and baseline interaction have same sign and opposite direction, in CS1 and CS2 this is classified as contradiction. Alternatively, the new interaction is flagged for further investigation in CS3, as the opposite direction may be possible under certain conditions. (3) : New interaction indicates that AMPK negatively and directly influences LKB1, and as such, it contradicts an interaction in the baseline graph in both sign and direction. It is also contradicted in locations of two entities. In CS1and CS2, this is classified as contradiction, however, in CS3, we classify it as a flagged interaction, assuming that it might fall into a correct interaction in another context. (4) : New interaction indicates that the presence of MYC increases the amount of TERT. In the baseline graph, there is neither an interaction nor a path between these two elements. In CS1 and CS3, this is classified as an extension, while in CS2, it is classified as flagged, since this may indicate a potential reader error when the new interaction is direct. (5) New interaction indicates that STAT3 indirectly activates IL6, however, there is no such interaction in the baseline graph, instead, there is a path with opposite direction, IL6 -> IL6R -> STAT3. In CS1 and CS3, this interaction is flagged for further investigation, while in CS2 it is classified as contradiction.

The magnitude of these changes varied across extraction systems (Figure 6B), reflecting differences in attribute availability. REACH and Llama 3 exhibited the strongest sensitivity to contextual attributes, indicating that these readers frequently extract richer contextual annotations. In contrast, INDRA showed minimal sensitivity beyond CA2, consistent with its limited contextual metadata, while GPT-4.1 demonstrated only small shifts across strategies, suggesting that optional attributes are extracted less consistently. Interactions derived from model B-relevant (R_B1-3_) and negative-control corpora (R_B*1,2_) were largely unaffected by attribute inclusion, indicating that contextual metadata was rarely present in those corpora.

These findings demonstrate that reconciliation outcomes are robust yet tunable. Attribute inclusion strategies allow users to calibrate reconciliation strictness to their modeling objectives without compromising structural integrity, using either broad pathway validation under CA1–CA2, or highly context-specific reconciliation under CA3–CA4.

### 2.9 Sensitivity to alternative mismatch semantics

Beyond attribute inclusion, interpretation of mismatches in directionality, connection type (direct vs. indirect), and paths may influence classification distributions. To evaluate the impact of semantic interpretation choices, we compared three classification schemes, CS1–CS3 (Methods), which differ in how direction mismatches and path-level inconsistencies are categorized (examples in Figure 6C).

Transitioning from the default scheme CS1 to the stricter mismatch interpretation in CS2 resulted in systematic increases in contradictions across all extraction systems (Figure 6A, column), reflecting the reclassification of path-aligned but attribute-inconsistent interactions as structural conflicts rather than ambiguous cases. Consequently, flagged interactions decreased for REACH, INDRA, and GPT-4.1, however they increased for Llama 3. Notably, Llama 3 is the only reader that triggers the reclassification of extensions into flagged cases, which occurs in VIOLIN’s decision tree specifically for direct interactions (Figure 2A). This aligns with observations that Llama 3 generates both a higher proportion of direct interactions in its output and the largest total number of extracted direct interactions relative to other readers (Figure S6). According to the decision tree, the increase in contradictions further indicates that many literature-derived interactions match existing model paths in source and target nodes but differ in sign or direction, suggesting opportunities to incorporate new feed-forward or feedback loops into the model.

By contrast, CS3 produces category distributions similar to CS1, as fewer interactions fall into cases treated differently by the two schemes. The relatively small changes in total category counts arise from opposing reclassifications that balance out at the aggregate level. Closer inspection shows that most prominent is the reclassification of sign contradictions from CS1 into flagged interactions under CS3, and these reclassified interactions may reveal novel information not captured in the baseline graphs.

Overall, these results indicate that the default scheme CS1 strikes a balance among the three schemes, enabling decisive classifications where possible while avoiding excessive assumptions about the contents of the interactions. Despite quantitative shifts, structural distributions remain stable across schemes, with extensions dominant and corroboration–contradiction balance changing predictably. This demonstrates that reconciliation semantics can be configured to match modeling objectives while preserving the underlying graph structure.

### 2.10 Modular integration with external knowledge constraints

To evaluate VIOLIN’s compatibility with external knowledge validation pipelines, we integrated database-backed filtering^30^ prior to reconciliation. Filtering removed interactions not supported by curated knowledge bases, reducing input noise and inconsistencies in grounding.

Following filtering, total interaction counts decreased substantially across extraction systems (Figure 7A). Disconnected full extensions were preferentially removed, while many corroborations and contradictions were retained. These results indicate that database filtering acts as a structural constraint that enriches for graph-consistent interactions while reducing speculative and erroneous expansion.

**Figure 7.**
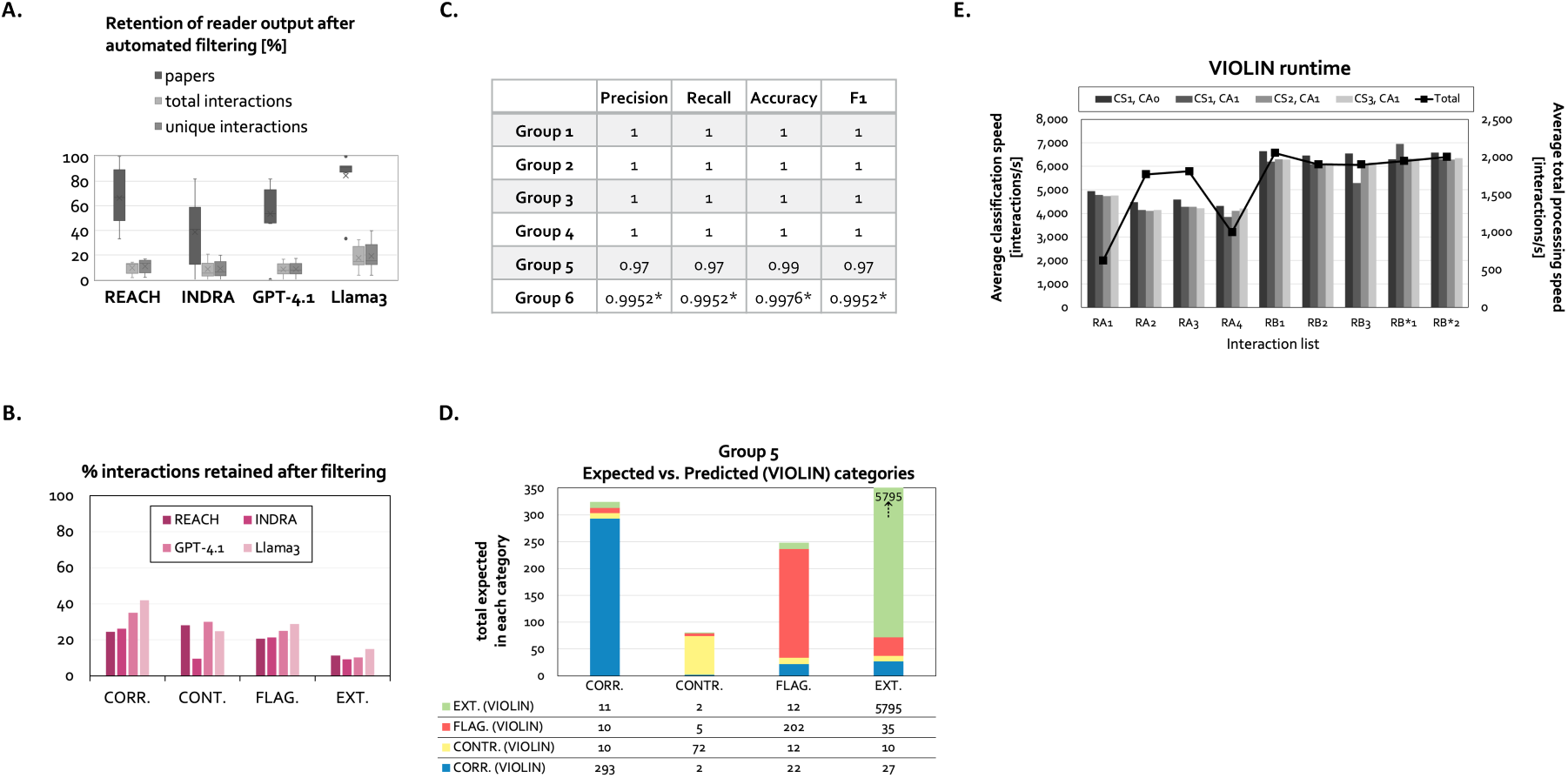
Integration with filtering, performance evaluation and runtime summary. **A.** The percent of papers, total interactions, and unique interactions retained after filtering, for each reader, across all corpora. **B.** The fraction (%) of interactions that are retained after filtering in each classification category in each reader output across all corpora except R_A1_. **C.** VIOLIN performance summary for each benchmark group (Methods), across all inputs. (*- only 4 interactions misclassified out of 831). **D.** Distribution of expected and predicted classifications in Group 5. For each category, the bar height represents total expected number of interactions in the category. Colors represented VIOLIN predictions. **E.** VIOLIN runtime. Bars- average classification speed 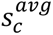 for each corpus, each classification scheme (CS1, CS2, CS3), and for CS1 comparing the classification strategy where none of the non-essential attributes are included (CA0) and the strategy where compartment information is included (CA1). Line-overall processing speed 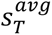 (loading inputs + classifying interactions + writing output to a file) averaged across all readers for CS1, CA0.

Retention patterns varied across readers, reflecting differences in grounding accuracy and identifier consistency (Figure 7B). Interactions that matched both source and target nodes of a baseline graph interaction - corroborations, contradictions, flagged, and internal extensions - were more likely to be preserved under filtering, whereas isolated extensions were more frequently removed (Figure S7A,B).

Comparison with semi-manual curation demonstrated that automated filtering is conservative, often removing interactions that human experts might retain for exploratory analysis (Figure S7C). VIOLIN’s modular design allows users to position filtering before or after reconciliation depending on modeling goals, providing a flexible mechanism for balancing precision and recall in knowledge integration workflows.

### 2.11 Algorithmic validation and agreement with expert curation

To validate algorithmic correctness and alignment with expert judgment, we evaluated VIOLIN across benchmark interaction lists of varying size and complexity (Methods). Controlled tests designed to confirm adherence to the decision tree (Groups 1-4) yielded perfect agreement with expected classifications, demonstrating correct implementation of match definitions (Figure 7C).

In larger benchmark comparisons against expert-curated classifications (Group 5), VIOLIN achieved high precision and recall (Figure 7C, D). Discrepancies stemmed from upstream entity grounding inconsistencies and not from reconciliation logic errors. In several instances, VIOLIN identified structural matches overlooked during manual curation, indicating that automated reconciliation can improve consistency and reduce human oversight variability.

When applied to filtered interaction lists with improved grounding quality (Group 6), classification agreement approached perfect precision and recall (Figure S8D). These results demonstrate that reconciliation accuracy is contingent on input grounding fidelity and confirm that VIOLIN’s classification logic operates as intended under properly standardized inputs. Together, these evaluations establish that VIOLIN reliably implements its formal decision rules and aligns closely with expert reasoning in structured graph comparison tasks.

### 2.12 Computational complexity and scalability

Runtime values indicate near-linear scaling with interaction list size (Table S2). Average classification time per interaction 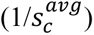 under the default scheme was approximately 0.18ms, with total processing time (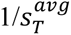 - includes loading, classification and writing time, Supplement Section 10) approximately 0.80ms per interaction, when run on a MacBook Pro with a 4.51 GHz Apple M4 Pro chip (Figure 7E). This performance represents multiple orders of magnitude acceleration relative to manual curation and remains substantially faster than upstream interaction extraction processes.

Runtime differences across corpora were primarily attributable to baseline graph size and path-finding rather than classification scheme selection. Consistent with expectations, interactions requiring path validation incurred greater computational cost than disconnected extensions. Overall, the computational complexity of VIOLIN’s classification algorithm ranges between O(*n* · *m*) and O(*n* · (*m* + (*m* + *e*) · log *m*)), where *n* is the number of extracted interactions, *m* is the number of nodes in the baseline graph, and *e* is the number of edges. This complexity arises from iterating over extracted interactions and performing path searches within the baseline graph.

These findings demonstrate that VIOLIN supports high-throughput reconciliation of large interaction graphs and is suitable for integration into automated knowledge graph updating pipelines. The combination of formalized decision rules, modular integration capacity, and scalable performance establishes VIOLIN as a practical infrastructure component for systematic literature-to-graph knowledge transfer.

## 3 Discussion

### Reconciliation as a structured graph problem

The central contribution of this work is the formalization of reconciliation between heterogeneous, literature-derived interaction graphs and structured baseline graphs as an attribute-aware graph comparison problem. By defining explicit match conditions for elements, directed signed interactions, and multi-edge paths, and by incorporating configurable attribute inclusion and mismatch semantics, VIOLIN provides a transparent and modular framework for structured knowledge integration. Rather than relying on ad hoc manual comparison or reader-specific heuristics, reconciliation is expressed as a formal, reproducible decision process that enables consistent comparison across heterogeneous extraction systems and supports scalable model updating workflows.

A key insight emerging from this work is that reconciliation outcomes are governed primarily by graph topology and attribute representation rather than corpus size alone. Extensions dominated classification results across all extraction regimes, indicating substantial structural incompleteness of curated baseline graphs relative to the broader literature landscape. Corroborations and contradictions were consistently present, revealing simultaneous reinforcement and opposition between curated knowledge and newly extracted evidence. Subtype analyses further demonstrated that path-level corroborations are prevalent, suggesting that literature frequently supports indirect regulatory structures encoded within baseline graphs. Conversely, sign and direction mismatches constituted the majority of contradictions, reflecting fundamental structural disagreements rather than superficial metadata discrepancies. Importantly, even under alternative classification schemes and attribute inclusion strategies, the overall structural distribution of categories remained stable — the dominance of extensions and the balance between corroborations and contradictions persisted, indicating that the framework captures intrinsic structural relationships between baseline graphs and extracted evidence rather than artifacts of parametric choices.

### Heterogeneous extraction and downstream stability

The inclusion of both rule-based NLP systems and LLM-based readers allowed evaluation of reconciliation under diverse extraction conditions. These systems differ substantially in extraction density, attribute completeness, and contextual richness, and we leveraged these differences as structured perturbations of the input graph space instead of considering them only as noise. LLM-derived graphs exhibited richer contextual attributes and more balanced contradiction subtypes, while traditional NLP-based systems demonstrated strengths in mechanistic annotation, particularly INDRA’s comprehensive mechanism coverage. Nevertheless, reconciliation remained stable across these differences, indicating that the framework is extraction-agnostic and robust to input variability. These findings are particularly relevant in the current landscape of rapidly evolving AI systems. As extraction technologies continue to diversify, downstream integration frameworks must remain adaptable to changing attribute richness and structural biases. By separating extraction from reconciliation and formalizing comparison at the graph level, VIOLIN provides a stable integration layer that can accommodate heterogeneous and evolving input sources without modification to its core logic.

### Configurability and modeling objectives

Another central feature of the framework is its configurability. Attribute inclusion strategies and alternative mismatch semantics allow users to tailor reconciliation criteria to specific modeling goals. For instance, inclusion of contextual attributes may be critical in highly specific cell-type models, whereas broader pathway validation may rely primarily on structural sign and direction agreement. Our results demonstrate that attribute inclusion and mismatch reinterpretation alter classification distributions in predictable and controlled ways. Reclassification proportions were generally modest relative to total interaction counts, suggesting that configurability enables fine-tuning of decision boundaries without significant changes to structural outcomes. This balance between flexibility and robustness is essential for practical model updating workflows. By explicitly encoding decision logic in configurable schemes, VIOLIN also makes interpretive assumptions transparent. This transparency is particularly important in interdisciplinary settings where domain experts may differ in how they interpret directional or contextual mismatches.

### Integration within modular curation pipelines

Integration with external database-supported filtering further demonstrates the modularity of the framework. Filtering preferentially removed disconnected extensions while preserving structurally consistent interactions, illustrating how reconciliation can be embedded within multi-stage knowledge validation pipelines. The comparison between automated filtering and semi-manual curation highlights trade-offs between conservativeness and exploratory expansion, reinforcing the importance of flexible workflow design. Given that reconciliation is independent of upstream extraction architecture and compatible with standardized graph formats, VIOLIN can function as an intermediate layer within automated literature-to-model pipelines. Extraction systems generate candidate interaction graphs, optional filtering modules constrain input based on curated databases, and reconciliation systematically classifies interactions relative to baseline graphs. This modular design supports scalable knowledge integration while preserving interpretability at each stage.

### Algorithmic correctness and expert alignment

Benchmark evaluations confirmed correct implementation of the formal decision tree and high agreement with expert curation under properly grounded inputs. Discrepancies were largely attributable to upstream grounding errors rather than classification logic. In several cases, automated reconciliation identified structural matches overlooked during manual inspection, demonstrating that VIOLIN can enhance consistency and reduce human oversight variability. These findings emphasize that reconciliation accuracy depends not only on algorithmic design but also on the quality of upstream extraction and entity grounding. Future integration of improved name recognition and grounding techniques that potentially also leverage advances in LLM-based normalization may further reduce classification discrepancies.

### Scalability and practical deployment

Scalability is essential for practical deployment in high-throughput literature integration workflows. Both theoretical complexity analysis and empirical runtime measurements demonstrate that VIOLIN scales approximately linearly with interaction list size and remains several orders of magnitude faster than manual curation. Path-based classifications incur higher computational cost than disconnected extensions, consistent with theoretical expectations, but remain computationally tractable even for large graphs. The combination of scalable performance, formalized decision logic, and modular compatibility positions VIOLIN as a practical infrastructure component for automated knowledge graph updating. As interaction databases expand and AI-based extraction systems continue to evolve, reconciliation frameworks must operate efficiently across increasingly large and diverse graph representations.

### Limitations and future directions

Several limitations warrant consideration. First, reconciliation accuracy is constrained by upstream entity grounding fidelity. Misidentified or inconsistently normalized entities may propagate to extension classifications or generate spurious mismatches. Improvements in entity normalization and identifier resolution will directly enhance reconciliation quality. Second, reconciliation currently operates on pairwise graph comparison and does not explicitly incorporate probabilistic confidence scores from extraction systems. Incorporating uncertainty estimates into classification decisions may provide additional nuance, particularly in distinguishing likely contradictions from low-confidence extraction artifacts. Finally, while the current evaluation focuses on biological signaling and regulatory networks, VIOLIN’s abstraction over directed, attributed graphs enables applicability to other domains where structured knowledge graphs evolve over time. Future work may explore extension to non-biological graph representations and integration with dynamic model updating and automated model assembly algorithms.

### Conclusion

We present VIOLIN as a configurable, attribute-aware reconciliation framework for integrating heterogeneous, literature-derived interaction graphs with curated baseline knowledge graphs. By formalizing reconciliation as a structured graph comparison problem and demonstrating robustness across extraction systems, attribute inclusion strategies, and semantic interpretations, we establish a scalable and modular infrastructure for systematic knowledge integration. As automated extraction technologies continue to expand the volume of structured interaction data, reconciliation frameworks such as VIOLIN will play a critical role in maintaining coherence between evolving literature and curated knowledge representations. The ability to quantify corroboration, contradiction, ambiguity, and expansion in a transparent and configurable manner provides a foundation for reproducible and scalable model updating in AI-assisted scientific discovery.

## 4 Methods

### 4.1 Formal match conditions and classification logic

VIOLIN evaluates each extracted interaction relative to a baseline graph using formally defined match conditions (Supplement Section 1.2). Two elements match if they share: 1- the same standardized identifier (HGNC symbol, name, or database ID), and 2- the same element type. We refer to these element attributes jointly as element’s identity. Two interactions satisfy the necessary interaction match condition if: 1- their source and target elements satisfy the element match condition, and 2- they have the same interaction sign. Attribute mismatches occur when corresponding attributes differ between matched interactions, excluding cases where one attribute value is empty. Provenance attributes are excluded from classification logic.

Each extracted interaction is classified relative to the baseline graph into one of four primary categories and multiple subcategories (formal definitions in Supplement Sections 1.3-1.6). Corroborations are interactions that satisfy necessary match conditions with a baseline interaction or path. Contradictions are interactions that match baseline elements but conflict in sign, direction, or other attributes with a baseline interaction. Extensions are interactions that introduce non-conflicting relationships that are not present in the baseline graph. Flagged are interactions that do not meet criteria for the other categories and require manual inspection. Path-based classification is performed using shortest-path searches within the baseline graph.

### 4.2 Generation of interaction lists

Interaction lists were generated using representative rule-based NLP systems (REACH, INDRA) and large language models (GPT-4.1, Llama 3). Workflows for finding literature corpora and extracting interactions are described in Supplement Sections 2 and 3 and outlined in Figure S3A. REACH^23^ machine reader relies on traditional rule-based NLP methods^23,58^. INDRA^28^ collects interactions from several NLP tools and databases and provides interaction belief scores based on their occurrence in literature and databases and on the probability of machine reading error. The INDRA database, with its millions of curated interactions, is a comprehensive and reliable source for mechanistic modeling. For REACH and INDRA, interaction extraction was most often performed directly from publication identifiers using their existing methods. For readers that required pre-fetching papers, we developed a structured workflow consisting of: 1- retrieval of full publication files; 2- conversion to machine-readable text format; 3- (LLMs only) few-shot prompting for structured interaction generation; 4-parsing of model output into tabular format. The prompt used for LLM extraction is provided in Figure S3C.

All extracted interactions were converted into BioRECIPE format using publicly available converter scripts that could also be accessed through the user-friendly interface (https://nmzlab.github.io/Tools-UI). Several examples of interactions with attributes are shown in Table S1. Search queries, paper counts, and extracted interaction counts are provided in Table S2. All files and scripts are available in the VIOLIN GitHub repository.

### 4.3 Baseline graph selection

For the majority of experiments, we used two baseline graphs with distinct biological context, structure, and size (Supplement Section 4). Baseline graph A is a directed, signed interaction network consisting of 179 nodes and 266 edges. The network includes protein–protein interactions, gene–mRNA–protein cascades, chemical regulations, and interactions involving biological processes. It contains 21 input nodes (nodes without regulators) and 9 output nodes (nodes without downstream targets). Mechanistic attributes are sparsely populated in this graph. Baseline graph B is a smaller directed interaction network containing 39 nodes and 70 edges, with 5 input nodes and 1 output node. This network predominantly represents protein–protein regulatory interactions and contains minimal chemical or transcriptional elements. Both graphs were represented using the BioRECIPE tabular format^48,50,51^, which encodes node identifiers, element types, interaction direction and sign, and optional contextual and mechanistic attributes. Visualizations of graphs A and B are shown in Figure S4A, B.

To further demonstrate compatibility with alternative knowledge sources, additional networks acquired from Reactome^1^ and Pathway Interaction Database^6,59^ (PID) were converted from BioPAX^60^ to BioRECIPE and used as baseline models when classifying one of the largest interaction lists (R_A2_). The summaries of all baseline graphs used for testing VIOLIN are provided in Figure S4C and all output files with classification results are available in the VIOLIN GitHub repository.

### 4.4 Attribute inclusion strategies (CA1–CA4)

In addition to necessary match conditions (element identity and interaction sign), VIOLIN allows optional inclusion of non-essential attributes in classification decisions. We evaluated four attribute inclusion strategies: CA1- compartment only; CA2- compartment and mechanism (CA1+mechanism); CA3-compartment, mechanism, and cell line (CA2+cell line); CA4- compartment, mechanism, cell line, and all the remaining contextual attributes (CA3+context). Under these strategies, mismatches in included attributes may result in reclassification (e.g., corroboration to contradiction or flagged) depending on the selected classification scheme. Attribute inclusion strategies enable users to calibrate reconciliation strictness based on modeling scope and contextual specificity requirements. Users can select any combination of non-essential attributes to be included in comparison through VIOLIN’s user interface.

### 4.5 Classification schemes (CS1–CS3)

VIOLIN can support different classification schemes that alter how directional and path-level mismatches are interpreted. These schemes modify decision boundaries while preserving necessary element and interaction match conditions. We implemented three schemes for evaluation. Differences between schemes are illustrated in Figure 2A (definitions in Supplement Sections 1.3-1.6). The default scheme that was used to collect most results is CS1. The unique characteristics of CS1 are: 1- direct interactions that align with a multi-edge path in the baseline graph are classified as extensions; 2- indirect interactions with attribute inconsistencies relative to a baseline path are classified as flagged. The second scheme, CS2 differs from CS1 in the following: 1- direct interactions that align with a baseline path are classified as flagged rather than extensions; 2- indirect interactions with mismatches in sign, direction, or non-essential attributes relative to a baseline path are classified as contradictions. The third scheme, CS3 differs from CS2 in the following: 1- direct interactions that align with an interaction in the baseline model in all attributes except direction are classified as contradictions rather than flagged; 2- when there is an additional attribute mismatch besides direction, direct interactions are classified as flagged instead of contradictions; other decision rules follow CS1. These schemes allow users to adjust semantic strictness in interpreting mismatches, enabling reconciliation behavior to be aligned with specific modeling objectives.

### 4.6 Integration with external filtering

To evaluate VIOLIN’s modular compatibility with external validation pipelines and literature-to-model workflows, we integrated VIOLIN with FLUTE^30^, a filtering tool that relies on interaction information from several databases. FLUTE retains interactions supported by at least one curated knowledge source. Filtered interaction lists were reconciled using VIOLIN to assess how database-driven constraints alter classification distributions (Supplement Section 8).

### 4.7 Benchmark construction

To validate implementation correctness and classification accuracy, we constructed six benchmark groups (detailed in Supplement Section 9, Figure S8). Groups 1–4 consist of synthetic and semi-synthetic interaction lists designed to test boundary cases of each classification subcategory and verify adherence to the formal decision tree. These tests confirmed correct implementation of element matching, interaction matching, path detection, and category assignment. Group 5 includes two large interaction lists manually classified by domain experts to evaluate agreement between algorithmic and expert reconciliation. Interaction lists were partitioned according to structural match conditions to enable targeted evaluation of extension, corroboration, and contradiction behavior. Group 6 consists of subsets of Group 5 interactions retained after database-supported filtering to assess classification accuracy under improved grounding conditions. Precision, recall, and F1 scores were computed for each benchmark group (Figure S8A).

### 4.8 Implementation

VIOLIN is implemented as a Python package with dependencies on Pandas, NumPy, and NetworkX. The framework operates on BioRECIPE-formatted^48,50,51^ interaction files and supports configurable classification schemes and attribute inclusion strategies. Path searches within baseline graphs are performed using Dijkstra’s algorithm as implemented in NetworkX. VIOLIN’s user interface is implemented as an interactive web application. The architecture of interface is based on a client-server model. Specifically, the frontend server is implemented with React, and backend server is built with FastAPI to serve the VIOLIN endpoints. Both frontend and backend are hosted on an AWS EC2 instance for online usage.

### 4.9 Data and code availability

All source code, baseline graphs, extracted interaction lists, benchmark datasets, and analysis scripts are publicly available in the VIOLIN GitHub repository (https://github.com/pitt-miskov-zivanov-lab/VIOLIN/). A Jupyter notebook is also included in the repository to facilitate reproducibility and reuse (https://github.com/pitt-miskov-zivanov-lab/VIOLIN/blob/master/examples/use_VIOLIN.ipynb). Scripts for format conversions to BioRECIPE are available, either as stand-alone (https://github.com/pitt-miskov-zivanov-lab/BioRECIPE) or through web-based user interface (https://nmzlab.github.io/Tools-UI).

## Acknowledgements

This work was funded by the DARPA Big Mechanism award W911NF-17-1-0135, the NSF EAGER award CCF-2324742, the NSF CAREER award CCF-2442884, and the NIH award R01LM014673.

## References

1. Milacic, M., Beavers, D., Conley, P., Gong, C., Gillespie, M., Griss, J., Haw, R., Jassal, B., Matthews, L., and May, B. (2024). The reactome pathway knowledgebase 2024. Nucleic acids research 52, D672–D678.

2. Kanehisa, M., Sato, Y., Kawashima, M., Furumichi, M., and Tanabe, M. (2016). KEGG as a reference resource for gene and protein annotation. Nucleic Acids Research 44, D457–D462. 10.1093/nar/gkv1070.

3. Ashburner, M., Ball, C.A., Blake, J.A., Botstein, D., Butler, H., Cherry, J.M., Davis, A.P., Dolinski, K., Dwight, S.S., and Eppig, J.T. (2000). Gene ontology: tool for the unification of biology. Nature genetics 25, 25–29.

4. UniProt: the Universal protein knowledgebase in 2025. (2025). Nucleic Acids Research 53, D609–D617.

5. Oughtred, R., Stark, C., Breitkreutz, B.-J., Rust, J., Boucher, L., Chang, C., Kolas, N., O’Donnell, L., Leung, G., McAdam, R., et al. (2019). The BioGRID interaction database: 2019 update. Nucleic Acids Research 47, D529–D541. 10.1093/nar/gky1079.

6. Rodchenkov, I., Babur, O., Luna, A., Aksoy, B.A., Wong, J.V., Fong, D., Franz, M., Siper, M.C., Cheung, M., and Wrana, M. (2020). Pathway Commons 2019 Update: integration, analysis and exploration of pathway data. Nucleic acids research 48, D489–D497.

7. Licata, L., Lo Surdo, P., Iannuccelli, M., Palma, A., Micarelli, E., Perfetto, L., Peluso, D., Calderone, A., Castagnoli, L., and Cesareni, G. (2020). SIGNOR 2.0, the SIGnaling Network Open Resource 2.0: 2019 update. Nucleic Acids Research 48, D504–D510. 10.1093/nar/gkz949.

8. Pratt, D., Chen, J., Welker, D., Kuentzer, J., Demchak, B., and Ideker, T. (2015). NDEx, the Network Data Exchange. Cell Systems 1, 302–305. 10.1016/j.cels.2015.10.001.

9. Malik-Sheriff, R.S., Glont, M., Nguyen, T.V., Tiwari, K., Roberts, M.G., Xavier, A., Vu, M.T., Men, J., Maire, M., and Kananathan, S. (2020). BioModels—15 years of sharing computational models in life science. Nucleic acids research 48, D407–D415.

10. Helikar, T., Kowal, B., and Rogers, J. (2013). A cell simulator platform: the cell collective. Clinical Pharmacology & Therapeutics 93, 393–395.

11. Fabregat, A., Sidiropoulos, K., Garapati, P., Gillespie, M., Hausmann, K., Haw, R., Jassal, B., Jupe, S., Korninger, F., McKay, S., et al. (2016). The Reactome pathway Knowledgebase. Nucleic Acids Research 44, D481–D487. 10.1093/nar/gkv1351.

12. Telmer, C.A., Sayed, K., Butchy, A.A., Bocan, K., Kaltenmeier, C., Lotze, M., and Miskov-Zivanov, N. (2021). Computational modeling of cell signaling and mutations in pancreatic cancer. preprint Systems Biology. 2021/06/09/. http://biorxiv.org/lookup/doi/10.1101/2021.06.08.447557.

13. Wang, Q., Miskov-Zivanov, N., Faeder, J.R., Lotze, M., and Clarke, E.M. (2016). Formal Modeling and Analysis of Pancreatic Cancer Microenvironment. 2016/09/21/23. (Springer), pp. 289–305.

14. Gómez Tejeda Zañudo, J., Mao, P., Alcon, C., Kowalski, K., Johnson, G.N., Xu, G., Baselga, J., Scaltriti, M., Letai, A., and Montero, J. (2021). Cell line–specific network models of ER+ breast cancer identify potential pi3kα inhibitor resistance mechanisms and drug combinations. Cancer research 81, 4603–4617.

15. Li, C., Wei, Y., and Lei, J. (2025). Quantitative cancer-immunity cycle modeling for predicting disease progression in advanced metastatic colorectal cancer. npj Systems Biology and Applications 11, 33.

16. Lavoie, H., Gagnon, J., and Therrien, M. (2020). ERK signalling: a master regulator of cell behaviour, life and fate. Nature reviews Molecular cell biology 21, 607–632.

17. Shin, S.-Y., Chew, N.J., Ghomlaghi, M., Chüeh, A.C., Jeong, Y., Nguyen, L.K., and Daly, R.J. (2024). Integrative Modeling of Signaling Network Dynamics Identifies Cell Type–Selective Therapeutic Strategies for FGFR4-Driven Cancers. Cancer Research 84, 3296–3309.

18. Blinov, M.L., Faeder, J.R., Goldstein, B., and Hlavacek, W.S. (2004). BioNetGen: software for rule-based modeling of signal transduction based on the interactions of molecular domains. Bioinformatics 20, 3289–3291.

19. Miskov-Zivanov, N., Turner, M.S., Kane, L.P., Morel, P.A., and Faeder, J.R. (2013). The Duration of T Cell Stimulation Is a Critical Determinant of Cell Fate and Plasticity. Science Signaling 6, ra97–ra97. 10.1126/scisignal.2004217.

20. Saez-Rodriguez, J., Simeoni, L., Lindquist, J.A., Hemenway, R., Bommhardt, U., Arndt, B., Haus, U.-U., Weismantel, R., Gilles, E.D., and Klamt, S. (2007). A logical model provides insights into T cell receptor signaling. PLoS computational biology 3, e163.

21. Zhang, Q., He, Y., Luo, N., Patel, S.J., Han, Y., Gao, R., Modak, M., Carotta, S., Haslinger, C., and Kind, D. (2019). Landscape and dynamics of single immune cells in hepatocellular carcinoma. Cell 179, 829–845. e820.

22. Li, X., and Xu, J.-X. (2016). A mathematical prognosis model for pancreatic cancer patients receiving immunotherapy. Journal of theoretical biology 406, 42–51.

23. Valenzuela-Escárcega, M.A., Hahn-Powell, G., Surdeanu, M., and Hicks, T. (2015). A Domain-independent Rule-based Framework for Event Extraction. Proceedings of ACL-IJCNLP 2015 System Demonstrations, 2015. (Association for Computational Linguistics and The Asian Federation of Natural Language Processing), pp. 127–132.

24. Ferguson, G., and Allen, J.F. (1998). TRIPs: an integrated intelligent problem-solving assistant. Proceedings of the fifteenth national/tenth conference on Artificial intelligence/Innovative applications of artificial intelligence. American Association for Artificial Intelligence.

25. Novichkova, S., Egorov, S., and Daraselia, N. (2003). MedScan, a natural language processing engine for MEDLINE abstracts. Bioinformatics 19, 1699–1706. 10.1093/bioinformatics/btg207.

26. Brown, T., Mann, B., Ryder, N., Subbiah, M., Kaplan, J.D., Dhariwal, P., Neelakantan, A., Shyam, P., Sastry, G., Askell, A., et al. (2020). Language Models are Few-Shot Learners. In H. Larochelle, M. Ranzato, R. Hadsell, M.F. Balcan, and H. Lin, eds.

27. Touvron, H., Martin, L., Stone, K., Albert, P., Almahairi, A., Babaei, Y., Bashlykov, N., Batra, S., Bhargava, P., Bhosale, S., et al. Llama 2: Open Foundation and Fine-Tuned Chat Models.

28. Gyori, B.M., Bachman, J.A., Subramanian, K., Muhlich, J.L., Galescu, L., and Sorger, P.K. (2017). From word models to executable models of signaling networks using automated assembly. Molecular Systems Biology 13, 954. 10.15252/msb.20177651.

29. Holtzapple, E., Cochran, B., and Miskov-Zivanov, N. (2021). Context-aware query design combines knowledge and data for efficient reading and reasoning. pp. 238–246.

30. Holtzapple, E., Telmer, C.A., and Miskov-Zivanov, N. (2020). FLUTE: Fast and reliable knowledge retrieval from biomedical literature. Database 2020. 10.1093/database/baaa056.

31. Hoyt, C.T., Domingo-Fernández, D., Aldisi, R., Xu, L., Kolpeja, K., Spalek, S., Wollert, E., Bachman, J., Gyori, B.M., Greene, P., and Hofmann-Apitius, M. (2019). Re-curation and rational enrichment of knowledge graphs in Biological Expression Language. Database: The Journal of Biological Databases and Curation 2019. 10.1093/database/baz068.

32. Ahmed, Y., Telmer, C., and Miskov-Zivanov, N. (2021). CLARINET: Efficient learning of dynamic network models from literature. Bioinformatics Advances. 10.1093/bioadv/vbab006.

33. Ahmed, Y., Telmer, C.A., Zhou, G., and Miskov-Zivanov, N. (2024). Context-aware knowledge selection and reliable model recommendation with ACCORDION. Frontiers in Systems Biology 4, 1308292.

34. Sayed, K., Bocan, K.N., and Miskov-Zivanov, N. (2018). Automated Extension of Cell Signaling Models with Genetic Algorithm. 2018 40th Annual International Conference of the IEEE Engineering in Medicine and Biology Society (EMBC), 2018/07//. (IEEE), pp. 5030–5033.

35. Butchy, A.A., Telmer, C.A., and Miskov-Zivanov, N. (2023). Automating Knowledge-Driven Model Recommendation: Methodology, Evaluation, and Key Challenges. arXiv [q-bio.MN].

36. Butchy, A.A., Telmer, C.A., and Miskov-Zivanov, N. (2021). FIDDLE: Efficient Assembly of Networks That Satisfy Desired Behavior. preprint In Review. 2021/06/02/.

37. Ahmed, Y., Butchy, A.A., Sayed, K., Telmer, C., and Miskov-Zivanov, N. (2021). New advances in the automation of context-aware information selection and guided model assembly. arXiv preprint arXiv:2110.10841.

38. Liang, K.-W., Wang, Q., Telmer, C., Ravichandran, D., Spirtes, P., and Miskov-Zivanov, N. (2017). Methods to Expand Cell Signaling Models Using Automated Reading and Model Checking. In Computational Methods in Systems Biology, J. Feret, and H. Koeppl, eds. (Springer International Publishing), pp. 145–159.

39. Holtzapple, E., Cochran, B., and Miskov-Zivanov, N. (2021). Automated verification, assembly, and extension of GBM stem cell network model with knowledge from literature and data. bioRxiv, 2021.2007.2004.451062. 10.1101/2021.07.04.451062.

40. Valenzuela-Escarcega, M.A., Babur, O., Hahn-Powell, G., Bell, D., Hicks, T., Noriega-Atala, E., Wang, X., Surdeanu, M., Demir, E., and Morrison, C.T. (2017). Large-scale automated reading with Reach discovers new cancer driving mechanisms. pp. 201–203.

41. Szklarczyk, D., Kirsch, R., Koutrouli, M., Nastou, K., Mehryary, F., Hachilif, R., Gable, A.L., Fang, T., Doncheva, N.T., and Pyysalo, S. (2023). The STRING database in 2023: protein–protein association networks and functional enrichment analyses for any sequenced genome of interest. Nucleic acids research 51, D638–D646.

42. Kuhn, M., Szklarczyk, D., Pletscher-Frankild, S., Blicher, T.H., Von Mering, C., Jensen, L.J., and Bork, P. (2014). STITCH 4: integration of protein–chemical interactions with user data. Nucleic acids research 42, D401–D407.

43. Gyori, B.M., Bachman, J.A., Subramanian, K., Muhlich, J.L., Galescu, L., and Sorger, P.K. (2017). From word models to executable models of signaling networks using automated assembly. Mol Syst Biol 13, 954. 10.15252/msb.20177651.

44. Agrawal, M., Hegselmann, S., Lang, H., Kim, Y., and Sontag, D. (2022). Large language models are few-shot clinical information extractors. arXiv preprint arXiv:2205.12689.

45. Dubey, A., Jauhri, A., Pandey, A., Kadian, A., Al-Dahle, A., Letman, A., Mathur, A., Schelten, A., Yang, A., and Fan, A. (2024). The llama 3 herd of models. arXiv preprint arXiv:2407.21783.

46. Cerami, E.G., Gross, B.E., Demir, E., Rodchenkov, I., Babur, Ö., Anwar, N., Schultz, N., Bader, G.D., and Sander, C. (2010). Pathway Commons, a web resource for biological pathway data. Nucleic acids research 39, D685–D690.

47. Hucka, M., Hoops, S., Keating, S., Le Novère, N., Sahle, S., and Wilkinson, D. (2008). Systems Biology Markup Language (SBML) Level 2: Structures and Facilities for Model Definitions. Nature Precedings. 10.1038/npre.2008.2715.1.

48. Holtzapple, E., Zhou, G., Luo, H., Tang, D., Arazkhani, N., Hansen, C., Telmer, C.A., and Miskov-Zivanov, N. (2024). The BioRECIPE knowledge representation format. ACS Synthetic Biology 13, 2621–2624.

49. Chandak, P., Huang, K., and Zitnik, M. (2023). Building a knowledge graph to enable precision medicine. Scientific Data 10, 67. 10.1038/s41597-023-01960-3.

50. BioRECIPE GitHub Repository. (2024). https://github.com/pitt-miskov-zivanov-lab/BioRECIPE/tree/main.

51. BioRECIPE Read the Docs. (2024). https://melody-biorecipe.readthedocs.io/en/latest/index.html.

52. Holtzapple, E., Cochran, B., and Miskov-Zivanov, N. (2021). Automated verification, assembly, and extension of GBM stem cell network model with knowledge from literature and data. preprint Systems Biology. 2021/07/05/. http://biorxiv.org/lookup/doi/10.1101/2021.07.04.451062.

53. Sayed, K., Telmer, C.A., Butchy, A.A., and Miskov-Zivanov, N. (2017). Recipes for translating big data machine reading to executable cellular signaling models. Machine Learning, Optimization, and Big Data.

54. Miskov-Zivanov, N. (2015). Automation of Biological Model Learning, Design and Analysis. Proceedings of the 25th edition on Great Lakes Symposium on VLSI, 2015/05/20/. (ACM), pp. 327–329.

55. Ahmed, Y., Telmer, C., and Miskov-Zivanov, N. (2022). Context-aware information selection and model recommendation with ACCORDION. bioRxiv, 2022.2001.2022.477231. 10.1101/2022.01.22.477231.

56. Ruscheinski, A., Wolpers, A., Henning, P., Wilsdorf, P., and Uhrmacher, A.M. (2025). SIMPROV: Provenance capturing for simulation studies. PLoS One 20, e0327607.

57. Uhrmacher, A.M., Frazier, P., Hähnle, R., Klügl, F., Lorig, F., Ludäscher, B., Nenzi, L., Ruiz-Martin, C., Rumpe, B., and Szabo, C. (2024). Context, composition, automation, and communication: The C2AC roadmap for modeling and simulation. ACM Transactions on Modeling and Computer Simulation 34, 1–51.

58. Valenzuela-Escárcega, M.A., Hahn-Powell, G., and Surdeanu, M. Odin’s Runes: A Rule Language for Information Extraction. Proceedings of the Tenth International Conference on Language Resource s and Evaluation (LREC’16), N. Calzolari, K. Choukri, T. Declerck, S. Goggi, M. Grobelnik, B. Maegaard, J. Mariani, H. Mazo, A. Moreno, J. Odijk, et al., eds. (European Language Resources Association (ELRA)), pp. 322–329.

59. Schaefer, C.F., Anthony, K., Krupa, S., Buchoff, J., Day, M., Hannay, T., and Buetow, K.H. (2008). PID: the Pathway Interaction Database. Nucleic Acids Research 37, D674–D679. 10.1093/nar/gkn653.

60. Demir, E., Cary, M.P., Paley, S., Fukuda, K., Lemer, C., Vastrik, I., Wu, G., D’Eustachio, P., Schaefer, C., Luciano, J., et al. (2010). The BioPAX community standard for pathway data sharing. Nature Biotechnology 28, 935–942. 10.1038/nbt.1666.

