## Supplementary materials for "VIOLIN: A modular framework for scalable reconciliation of heterogeneous interaction graphs"

### Supplementary Material

#### 1 Formal definitions and notation

This section provides the formal definitions underlying VIOLIN's reconciliation algorithm.

##### 1.1 Element, interaction, and path definitions

Several definitions in this sub-section are adopted from the BioRECIPE documentation[1,2] and are included here to provide notation context for new definitions. Table S1 includes five examples of element and interaction attribute values. More examples can also be found in GitHub repositories for BioRECIPE[3] and VIOLIN[4].

*Definition 1.1.* An ***element*** $v = v(\mathbf{a}^v)$, is defined by its name ($a^{name}$), type ($a^{type}$), and a unique database identifier (ID) ($a^{database}$ and $a^{ID}$) or the gene symbol from the HGNC database ($a^{HGNCsymbol}$). These five attributes are referred to as *essential element attributes*. Other element attributes are the element subtype $a^{subtype}$, the cellular compartment $a^{compartment}$ and the compartment's unique ID $a^{compartmentID}$. Element attributes are written as a vector:

$$\mathbf{a}^v = \left(a^{name}, a^{type}, a^{subtype}, a^{HGNCsymbol}, a^{database}, a^{ID}, a^{compartment}, a^{compartmentID}\right).$$

*Definition 1.2.* A ***directed signed interaction*** $e = e(v_s, v_t, \mathbf{a}^e)$ is defined with its source element $v_s$, target element $v_t$, and a vector of attributes $\mathbf{a}^e$. The *essential interaction attribute* is the interaction sign $a^{sign}$. The direction of an interaction is always implicitly defined with its source and target nodes, and therefore, not explicitly listed among its attributes. Other interaction attributes that further define the **influence** are the interaction connection type $a^{connectiontype}$ indicating whether the influence is direct or indirect, the mechanism of the interaction $a^{mechanism}$, and the molecular site of the interaction $a^{site}$. The **context** attributes include cell line $a^{cellline}$, cell type $a^{celltype}$, tissue type $a^{tissuetype}$, and organism $a^{organism}$ where the interaction occurs. The **provenance** attributes include the belief or confidence score for the interaction $a^{score}$, the source from which the interaction is obtained $a^{source}$, the statements $a^{statements}$ and IDs of papers $a^{paperIDs}$ where the interaction is mentioned. These attributes together form the interaction attribute vector:

$$\mathbf{a}^e = (a^{sign}, a^{connectiontype}, a^{mechanism}, a^{site}, a^{cellline}, a^{celltype}, a^{tissuetype}, a^{organism}, a^{score}, a^{source}, a^{statements}, a^{paperIDs}).$$

*Definition 1.3.* A ***path*** in a model is formed from $n > 1$ connected edges: $p\left(v_{s_p}, v_{t_p}, a^{sign_p}\right) = (e(v_{s_p} = v_{k_1}, v_{k_2}, \mathbf{a}^e_{k_1}),\ e(v_{k_2}, v_{k_3}, \mathbf{a}^e_{k_2}), \ldots, e(v_{k_n}, v_{k_{n+1}} = v_{t_p}, \mathbf{a}^e_{k_n}))$. The direction of the path is implicitly defined with the source node $v_{s_p}$ and target node $v_{t_p}$. The regulation sign $a^{sign_p}$ is considered positive when the number of negative signs in the set $\{a_{k_1}^{sign}, a_{k_2}^{sign}, .., a_{k_n}^{sign}\}$ is even, and negative when this number is odd.

*Definition 1.4.* A directed signed interaction $e = e(v_s, v_t, \mathbf{a}^e)$ where $v_s \equiv v_t$ is called ***direct self-regulation***.

*Definition 1.5.* A path $p(v_{s_p}, v_{s_p}, a^{sign_p})$ where the path source is also the path target, $v_{s_p} \equiv v_{s_p}$, is called a ***feedback loop***. If $a^{sign_p}$="positive", it is a positive feedback loop and if $a^{sign_p}$="negative", it is a negative feedback loop.

#### 1.2 Match definitions

*Definition 2.1.* Two elements $v = v(\mathbf{a}^v)$ and $v' = v'(\mathbf{a}^{v'})$ satisfy the ***necessary element match condition*** if they have at least one of the following, the same HGNC symbol ($a^{HGNCsymbol} = a^{HGNCsymbol'}$) or the same name ($a^{name} = a^{name'}$) or the same unique database ID ($a^{database} = a^{database'}$ and $a^{ID} = a^{ID'}$), and they also have the same element type ($a^{type} = a^{type'}$).

*Definition 2.2.* Two interactions, $e(v_s, v_t, \mathbf{a}^e)$ and $e'(v'_s, v'_t, \mathbf{a}^{e'})$, satisfy the ***necessary interaction match condition*** if elements $v_s$ and $v_t$ satisfy the necessary condition to match elements $v'_s$ and $v'_t$, respectively, and the interaction signs are same ($a^{sign} = a^{sign'}$).

*Definition 2.3.* If the corresponding attributes in two interactions are different (e.g., $a^{location} \neq a^{location'}$), we refer to this as ***attribute mismatch***. We note here that, in the case when the attribute value in one of the two interactions is "empty", this will not be considered as an attribute mismatch.

*Definition 2.4.* An interaction $e'(v'_s, v'_t, \mathbf{a}^{e'})$ satisfies the ***necessary path match condition*** to match a path in a model, if $a^{connectiontype'}$ ="indirect", there exists a path $p\left(v_{s_p}, v_{t_p}, a^{sign_p}\right)$ in the model, nodes $v'_s$ and $v'_t$ satisfy the necessary conditions to match two nodes $v_{s_p}$ and $v_{t_p}$, and the sign of the interaction matches the sign of the path $a^{sign'} = a^{sign_p}$.

#### 1.3 Corroboration definitions

The definitions in sections 1.3-1.6 assume the default classification scheme CS1, and classification attribute choice strategy CA1. When additional conditions are included for other schemes (CS2, CS3) or strategies (CA2, CA3, CA4), a NOTE following the definition describes these differences.

*Definition 3.1.* An interaction $e'(v'_s, v'_t, \mathbf{a}^{e'})$ is classified as ***weak corroboration*** of a model interaction $e(v_s, v_t, \mathbf{a}^e)$, if $e$ and $e'$ satisfy the necessary interaction match condition and one of the following conditions is satisfied:

i) ***empty attribute***: there is no mismatch between corresponding attributes in $\mathbf{a}^e$ and $\mathbf{a}^{e'}$, and at least one of the non-essential attributes in $\mathbf{a}^{e'}$ is "empty".

ii) ***indirect interaction***: there is no mismatch between corresponding attributes in $\mathbf{a}^e$ and $\mathbf{a}^{e'}$, except $e'$ is an indirect interaction, $a^{connectiontype'}$ = "indirect" and $e$ is a direct interaction, $a^{connectiontype}$="direct".

*Definition 3.2.* An interaction $e'(v'_s, v'_t, \mathbf{a}^{e'})$ with $a^{connectiontype'}$ = "indirect" is classified as ***path corroboration*** if there is no such interaction $e(v_s, v_t, \mathbf{a}^e)$ in the model that satisfies necessary element match conditions with $e'$, and instead $e'$ satisfies the necessary condition to match a path $p\left(v_{s_p}, v_{t_p}, a^{sign_p}\right)$ in the model.

*Definition 3.3.* An interaction $e'(v'_s, v'_t, \mathbf{a}^{e'})$ is classified as ***strong corroboration*** of a model interaction $e(v_s, v_t, \mathbf{a}^e)$, if $e$ and $e'$ satisfy the necessary interaction match condition, $a^{connectiontype'}$ = $a^{connectiontype}$ and the $a^{compartment'}$ and $a^{compartmentID'}$ attribute values of both $v'_s$ and $v'_t$ match the values of corresponding attributes in $v_s$ and $v_t$, respectively (NOTE: in the other three classification strategies, additional attributes are included in the condition, specifically, the mechanism attribute for CA2, mechanism and cell line attributes for CA3, and mechanism and all context attributes for CA4).

*Definition 3.4.* An interaction $e'(v'_s, v'_t, \mathbf{a}^{e'})$ is classified as ***specification*** if there exists a model interaction $e(v_s, v_t, \mathbf{a}^e)$, such that $e$ and $e'$ satisfy the necessary interaction match condition, and either

i) all remaining attributes match such that there is at least one "empty" model interaction attribute and a non-"empty" corresponding interaction attribute, or

ii) $a^{connectiontype'}$="direct" and $a^{connectiontype}$="indirect" and all the remaining attributes match.

##### 1.4 Contradiction definitions

*Definition 4.1.* An interaction $e'(v'_s, v'_t, \mathbf{a}^{e'})$ is classified as ***contradiction*** if there exists a baseline model interaction $e(v_s, v_t, \mathbf{a}^e)$ such that:

i) ***direction contradiction***: $v'_t$ satisfies the necessary condition to match $v_s$, $v'_s$ satisfies the necessary condition to match $v_t$ , and either the model interaction is indirect

($a^{connectiontype}$=”indirect”), or the model interaction is direct ($a^{connectiontype}$=”direct”) and there is a mismatch between $e'$ and $e$ in at least one non-essential attribute. (NOTE: in the CS3 scheme, this is considered a flagged interaction, if there is also a sign mismatch, Figure 3).

ii) ***sign contradiction***: $v'_s$ satisfies the necessary condition to match $v_s$, $v'_t$ satisfies the necessary condition to match $v_t$, and there is a mismatch in signs between $e$ and $e'$, that is, $a^{sign} \neq a^{sign'}$. (NOTE: in the CS3 scheme, when $a^{connectiontype'}$ =”indirect” and $a^{connectiontype}$=”direct” this is considered a flagged interaction, Figure 3).

iii) ***attribute conflict***: $e$ and $e'$ satisfy the necessary match conditions, and there is a mismatch between $e$ and $e'$ in at least one non-essential attribute other than $a^{connectiontype'}$.

The reasoning behind the connection type requirement in the direction contradiction definition is that an indirect model interaction may be a “placeholder” in the absence of more detailed knowledge, and the new interaction can dispute this previously incomplete knowledge. On the other hand, when the model interaction is direct, and there is a contradiction with the new interaction in direction and other attributes (e.g., location), this can indicate a potential error in the interaction extraction tool or an incorrect model interaction.

##### 1.5 Extension definitions

*Definition 5.1.* An interaction $e'(v'_s, v'_t, \mathbf{a}^{e'})$ is classified as ***extension*** in the following cases:

iii) ***internal extension***: if there exist two different model elements $v_i$ and $v_j$ such that $v'_s$ and $v'_t$ satisfy at least the necessary condition to match $v_i$ and $v_j$, respectively, there is no interaction $e(v_i, v_j)$ or interaction $e(v_j, v_i)$ in the model, and either $a^{connectiontype'}$ =”direct”, or $a^{connectiontype'}$=”indirect” and there is no path $p(v_i, v_j)$ in the model. (NOTE: in the CS2 scheme, when $a^{connectiontype'}$=”direct”, this is considered a flagged interaction, Figure 3).

iv) ***hanging extension***: if either $v'_s$ or $v'_t$ (and not both) matches a model element.

v) ***full extension***: there are no baseline model elements that match $v'_s$ and $v'_t$.

##### 1.6 Flagged definitions

There are some cases where a combination of matched, mismatched, and “empty” attributes requires further manual inspection before it can be classified, and therefore, these interactions are flagged by VIOLIN.

*Definition 6.1.* An interaction $e'(v'_s, v'_t, \mathbf{a}^{e'})$ is classified as ***flagged*** in the following cases:

i) ***direction mismatch***: if there exists a model interaction $e(v_s, v_t, \mathbf{a}^e)$ with $a^{connectiontype}$="direct", $v'_s$ and $v'_t$ satisfy the necessary condition to match $v_t$ and $v_s$, respectively, and there is no mismatch in other element attributes and no mismatch in context attributes ("empty" attributes are allowed). (NOTE: in the CS3 scheme, this is in some instances considered a direction contradiction, Figure 3).

ii) ***path mismatch***: if $a^{connectiontype'}$="indirect", nodes $v'_s$ and $v'_t$ satisfy the necessary condition to match two nodes in the baseline model, $v_i$ and $v_j$, respectively, there is no interaction $e(v_i, v_j)$ or interaction $e(v_j, v_i)$ in the model, and instead there is either a path $p(v_i, v_j, a^{sign_p})$ where $a^{sign'} \neq a^{sign_p}$ or a mismatch in any of the non-essential attributes of the source and target elements, or there is a path $p(v_j, v_i, a^{sign_p})$. (NOTE: in the CS2 scheme, these are considered contradictions, Figure 3).

iii) ***self-regulation***: if $e'$ is a self-regulation ($v_s \equiv v_t$) and there exists a model element $v_i$ such that $v'_s$ and $v'_t$ satisfy the necessary condition to match $v_i$.

The self-regulation flagged interactions may indeed be self-regulations, but more often the self-regulation is a result of grounding in the interaction extraction tool, and the level of detail and abstraction in the baseline model, and therefore, we flag them. Grounding is an extraction step that uses public databases [5,6] to pair a standard identifier with the common name used in the literature text, though it can occasionally result in error. For example, in the literature, Caspase-3 and Caspase-8 interact with each other as separate entities, while in the melanoma model that we use as one of our case studies, all the caspases are grouped in a single variable. Thus, the interaction between Caspase-3 and Caspase-8 is classified as self-regulation, and the user can then decide how to use the interaction. Self-regulation interactions may also result from the qualifier such as "mutated" or "phosphorylated" being ignored by the reader. For example, the statement "R2834H DP decreases phosphorylation of DP compared with WT" means that the mutated protein is not phosphorylated and therefore decreases the phosphorylation compared to wild type (WT), the extracted event is "DP negatively regulates DP," ignoring the important distinction of the mutated form.

#### 2 Corpus construction and query design

To generate heterogeneous interaction inputs for reconciliation analysis, we constructed nine literature corpora. These corpora were designed to vary in size, query specificity, contextual alignment with baseline graphs and retrieval strategy with varying degrees of topical alignment to the baseline graphs.

Corpora were assembled using PubMed Central[7] and REACH Explorer[8] queries designed to produce: 1- broad, context-aligned literature set ($S_{A1}$); 2- protein-centered retrieval strategy ($S_{A2}$); 3- targeted pathway-focused literature sets for baseline graphs A and B ($S_{A3}$, $S_{A4}$, $S_{B1}$ - $S_{B3}$); 4- intentionally weakly related ("negative") literature sets ($S_{B*1}$, $S_{B*2}$). Query details are provided in Table S2. Relevant corpora were designed to align with the biological contexts represented by the baseline graphs. Negative corpora included terms unrelated to one baseline graph to evaluate reconciliation robustness under irrelevant input conditions. This variation allowed evaluation of VIOLIN under diverse input conditions.

#### 3 Reader extraction behavior

The steps that we used for generating interaction lists are outlined in Figure S3A and the features of the four tools relevant to interaction extraction are summarized in Figure S3B. The prompt used with LLMs is included in Figure S3C. The summary of paper corpus sizes and corresponding interactions list sizes are provided in Table S2. An average runtime per paper across all interaction lists (Figure S3B) indicates that INDRA outperforms the other tools, while GPT-4.1 is the slowest. This difference in speed is expected, given that INDRA draws its output in part from existing databases while GPT-4.1 is based on a very large model. Further, for the largest paper corpus, $S_{A1}$, we limited analysis to REACH and INDRA only, as applying LLM-based readers to this corpus was computationally impractical due to extended runtimes.

#### 4 Baseline knowledge graphs - context and source

Baseline graph A (melanoma signaling network) was created based on biological pathways in the melanoma SkMel-133 cell line from Korkut et al [9], where the authors performed DNA, RNA, and protein analysis on primary and metastatic melanomas from 331 patients. Their investigation focused on four specific genomic subtypes: mutant BRAF, mutant RAS, mutant NF1, and triple wild type. The goal of both the wet lab experiments and the model analysis was to investigate potential immunotherapies. Additional subtypes exist that may benefit from MAPK and RTK inhibition therapies[9]. Graph topology is relatively dense compared to graph B, with multiple convergent signaling cascades and cross-pathway interactions (Figure S4A).

Baseline graph B (naïve T cell differentiation network) was created based on the published network[10] that captures the circuitry that controls the differentiation of naïve T cells into regulatory (Treg) and helper (Th) cells. Previously published model analysis[10] focused on the early steps of T cell activation, and the suppression or generation of regulatory T cells (Treg), connecting the Akt-mTOR pathway to Treg development and the T cell receptor (TCR) pathways to activation and inhibition of Foxp3, leading to different T cell fates. In contrast to the melanoma SkMel-133 cell network, the T cell network includes almost exclusively protein-protein interactions, with very few chemical and gene nodes. Compared to graph A, this network is smaller and topologically simpler, providing a contrasting structural environment for reconciliation (Figure S4B).

We also classified interaction list $R_{A2}$ with respect to several other models. The summaries of all baseline graphs used for testing VIOLIN are provided in Figure S4C. Results for these additional models are available on GitHub.

#### 5 Detailed classification subcategory distribution

We investigated classification subtype distributions across all readers and all corpora. The distributions and total numbers for each reader in each category are shown in Figure S5.

#### 6 Attribute coverage by readers

In addition to attribute coverage for each reader across all corpora (Figure 3B), we also summarized attribute coverage using two heatmaps (Figure S6). The first heatmap (Figure S6A) shows relative attribute coverage within each extracted interaction list, for each reader and each corpus. The second heatmap (Figure S6B) shows counts of interactions that have non-empty or specific values for mechanism and connection type attributes, colored relative to the highest count for each corpus.

Although finding new direct interactions and including the information about their mechanism is critical for mechanistic modeling, the results suggest that traditional machine readers struggle to determine whether an interaction is direct, and therefore, default to assigning "indirect" to the connection type attribute, while LLM tools are more successful in this task. On the other hand, INDRA assigns values to the mechanism attribute in all extracted interactions. However, given the small number of interactions INDRA finds, Llama 3 extracts the largest number of interactions with Mechanism attribute value in most corpora.

#### 7 Sensitivity to attributes within classification criteria

The classification results highlight the importance of carefully selecting the most relevant literature for modeling and offer insights into the most useful approach for information extraction based on the classification category of interest.

Non-essential attributes may skew the classification outcome. For instance, the baseline graph for SkMel133 cell line includes interaction 'LKB1' activates 'AMP-activated protein kinase (AMPK)' and an extracted interaction states that 'AMPK' is positively regulated by 'LKB1', with the literature evidence linking it to NCI-H460, a Non-Small Cell Lung Cancer (NSCLC) cell line. When only necessary match attributes and the compartment attribute are used (CA1 strategy), the extracted interaction is classified as strong corroboration. When the cell line attribute is included in classification criteria (CA3), this interaction is classified as contradiction due to the conflicting cell line attribute.

When changing from CA1 to CA4, corroborations are reclassified to contradictions and flagged for REACH (57% of corroborations; 1.4% of all interactions) and Llama 3 (40% of corroborations, 2.9% of all interactions), and few being reclassified to internal extensions (Figure 6B). While adding Mechanism to classification strategy in CA2 had very little influence on classification outcomes overall, it was the only change in criteria that affected INDRA output (6 interactions total), as it didn't further change in CA3 and CA4. This is expected, as INDRA does not provide values for contextual attributes such as cell line, cell type, tissue type, or organism (Figure 3B). Interestingly, GPT-4.1 output is not affected by the inclusion of the mechanism attribute in the classification criteria and instead shows its greatest sensitivity to contextual criteria. Even so, only 17 interactions are reclassified in its output when changing from CA1 to CA4 across $S_{A2}$-$S_{B*2}$ corpora, compared to 164 for REACH and 154 for Llama 3.

#### 8 Database-supported reader output retention

Figure S7 provides retention values for each subcategory and each reader across corpora. Summarized across $R_{A2}$-$R_{B*1}$, LLM output has higher retention rate than NLP in most subcategories (Figure S7A). The outliers with 100% retention are cases where only one interaction was extracted in the subcategory and retained, and therefore, using the largest corpus is a better indicator of the quality of reader output. For $R_{A1}$, REACH and INDRA have similar retention rate in most subcategories (Figure S7B). Consistently for all corpora, INDRA did not provide strong corroborations and attribute conflicts due to its lower attribute coverage.

Internal extensions are more likely to be supported by interaction databases, as they are the only extension subcategory where interactions match both model elements. This is confirmed by results (Figure S7A,B), which show that internal extensions are filtered out significantly less than full and hanging extensions.

We note that the number of interactions considered within classification subcategories can vary several orders of magnitude (Figure S7A,B - line), and only ~10 total strong corroborations are retained across all corpora, versus ~10,000 retained extensions for $R_{A1}$. Lower than expected retention of strong corroborations is likely due to the filtering tool removing all protein family names (e.g., AKT) and keeping only specific proteins (e.g., AKT1, AKT2), which results in filtering out even some interactions that VIOLIN classified as strong corroborations. Thus, VIOLIN may be less restrictive in some cases, considering a match as long as the same element ID and element type are used.

#### 9 Benchmark details and evaluation results

We created several groups of manually curated benchmark interaction lists. All interaction lists are in the form of Excel files and included in the VIOLIN GitHub repository (https://github.com/pitt-miskov-zivanov-lab/VIOLIN/tree/master/eval). The summary of lists is provided in Figure S8A. All scripts and the spreadsheets with results are also available on GitHub.

**Group 1** has 14 fully populated interactions, with one interaction in each subcategory with respect to model A. These interactions were created manually to test the correctness of the algorithm implementation - whether VIOLIN follows the decision tree outlined in Figure 2A.

**Group 2** includes two interactions lists with interactions from either model A (266 interactions) or model B (70 interactions). These two benchmark lists were created with a script that converts models to corresponding interaction lists and are used to test the accuracy of classifying into the corroboration category. The translator is available as part of the UI, under BioRECIPE (https://nmzlab.github.io/Tools-UI), as well as a separate script code on GitHub (https://github.com/pitt-miskov-zivanov-lab/BioRECIPE/tree/main/translators/within_biorecipe).

**Group 3** includes two interaction lists that were created by adding 10 contradictions for models A and B to the two corresponding lists in Group 2.

**Group 4** includes two interaction lists that were created by adding 10 randomly selected and manually classified interactions from REACH and INDRA outputs to interaction lists in Group 2.

**Group 5** was created to evaluate the effectiveness and human expert alignment of the algorithm on a larger set. Since manual curation of interaction lists is slow, we chose two interaction lists from REACH output, one from each set A and B, $R_{A2}$ and $R_{B2}$ (sizes listed in Figure S4). For these two lists, we created new sub-lists $M_0$, $M_1$, $M_2$, and $M_3$ (Figure S8B). The two $M_0$ lists were created by removing interactions where the regulator or regulated element name either contains multiple entities, begins with a number, or consists solely of special characters (e.g., #, /, ~, $, etc.). $R_{A2}$-$M_0$ list has 5,274 unique interactions, $R_{B2}$-$M_0$ list has 411 unique interactions. $M_1$ was formed from $M_0$ by keeping only the interactions that do not have any element matches with the model. $M_2$ consists of interactions where exactly one element matches a model element. The remaining interactions that match the model in both elements formed list $M_3$. By construction, all interactions in lists $M_1$ are full extensions and all interactions in lists $M_2$ are hanging extensions, with respect to their corresponding model. Interactions in $M_3$ lists are either corroborations, contradictions, flagged, or internal extensions, with respect to their corresponding model. The two $M_3$ lists were manually classified to serve as a benchmark for evaluating VIOLIN's classification.

**Group 6** includes only those interactions from Group 5 that were retained after filtering with FLUTE, reducing lists $R_{A2}$-$M_0$ and $R_{B2}$-$M_0$ to 870 and 37 interactions, respectively (Figure S8C).

As expected, when compared to manual curation, VIOLIN had 100% correct performance on all benchmarks in Groups 1-4 (Figure S8A). In Group 5, VIOLIN has 100% precision and recall for benchmark $M_1$ in $R_{B2}$ and has >97.5% recall for $M_1$ in $R_{A2}$ (3 out of 2,250 interactions misclassified). Inspection of misclassified interactions confirms again that VIOLIN correctly implements the decision tree in Figure 2A and that the errors stem from the REACH entity recognition and grounding – while an expert is able to recognize nuances in element and interaction attributes, VIOLIN's string processing was not created to handle errors and assumes correctly grounded entities. In our future work, we will explore methods to improve name recognition in VIOLIN, possibly using LLMs. To confirm that VIOLIN can indeed correctly classify interactions that are properly grounded, using Group 6 benchmarks we examined classification accuracy of the interactions that were retained after filtering with FLUTE. Since these interactions are highly supported by databases, they are less likely to have grounding errors, and therefore, should be accurately classified by VIOLIN.

Notably, upon initial inspection of misclassified interactions, we found that VIOLIN corrected the human curator in several instances – VIOLIN was able to identify a matching interaction or pathway in the model that the human curator missed. These results highlight VIOLIN's ability to both correct human errors and significantly accelerate the curation process beyond what is possible with manual methods.

#### 10 Extended runtime and complexity analysis

The total VIOLIN runtime $T_T$ is calculated as follows:

$$T_T = T_L + T_C + T_W$$

$T_L$ – loading time – the time necessary to read the baseline model file (or interaction list) and the new interaction list file

$T_C$ – classification time – the time necessary to process an interaction list with VIOLIN

$T_W$ – write time – the time necessary to create output files for the four categories

We also computed the average classification speed as:

$$s_C^{avg} = avg(s_{C,REACH}, s_{C,INDRA}, s_{C,ChatGPT3.5}, s_{C,Llama2})$$

where the classification speed for each reader output is calculated as a ratio between the number of interactions that VIOLIN treats as unique in the reader output and the classification time for those interactions:

$$s_{c,reader} = \frac{N_{VIOLIN\ unique,reader}}{T_{C,reader}}$$

The average total processing speed is calculated as:

$$s_T^{avg} = avg(s_{T,REACH}, s_{T,INDRA}, s_{T,ChatGPT3.5}, s_{T,Llama2})$$

where the total processing speed for each reader output is calculated as a ratio between the total number of interactions output by the reader and the total time needed for VIOLIN to load input files, classify interactions, and write output files for that reader output:

$$s_{T,reader} = \frac{N_{T,reader}}{T_{T,reader}}$$

**Algorithm complexity analysis**. VIOLIN takes two inputs: an interaction list containing $n$ interactions and a model whose graph topology consists of $m$ nodes and $e$ edges. VIOLIN processes one interaction per iteration, where finding matching nodes in the baseline graph is approximately $O(m)$. In the worst case, a search for a path between matched nodes is necessary, and VIOLIN then invokes a path-finding Dijkstra's algorithm, contributing $O((m+e)\cdot \log m)$ per iteration. Therefore, the total time complexity is bounded between $O(n\cdot m)$ and $O(n(m+(m+e)\cdot \log m))$. We also note that string matching for entities may vary due to different algorithms, and here we treat it as $O(1)$ and exclude it from this analysis.

#### References


1. Holtzapple, E., Luo, H., Tang, D., Zhou, G., Arazkhani, N., Hansen, C., Telmer, C.A., and Miskov-Zivanov, N. (2024). The BioRECIPE Knowledge Representation Format. bioRxiv, 2024.2002. 2012.579694.

2. BioRECIPE: Biological system Representation for Evaluation, Curation, Interoperability, Preserving, and Execution. (2023). https://melody-biorecipe.readthedocs.io/en/latest/index.html.

3. BioRECIPE GitHub Repository. (2024). https://github.com/pitt-miskov-zivanov-lab/BioRECIPE/tree/main.

4. VIOLIN GitHub Repository. (2024). https://github.com/pitt-miskov-zivanov-lab/VIOLIN.

5. Ashburner, M., Ball, C.A., Blake, J.A., Botstein, D., Butler, H., Cherry, J.M., Davis, A.P., Dolinski, K., Dwight, S.S., Eppig, J.T., et al. (2000). Gene Ontology: tool for the unification of biology. Nature Genetics *25*, 25-29. 10.1038/75556.

6. Consortium, T.U. (2018). UniProt: a worldwide hub of protein knowledge. Nucleic Acids Research *47*, D506-D515. 10.1093/nar/gky1049.

7. PubMed Central. (2005). Reference Reviews *19*, 37-38. doi:10.1108/09504120510587797.

8. Valenzuela-Escárcega, M.A., Hahn-Powell, G., Surdeanu, M., and Hicks, T. (2015). A Domain-independent Rule-based Framework for Event Extraction. held in Stroudsburg, PA, USA, (Association for Computational Linguistics and The Asian Federation of Natural Language Processing), pp. 127-132.

9. Akbani, R., Akdemir, Kadir C., Aksoy, B.A., Albert, M., Ally, A., Amin, Samirkumar B., Arachchi, H., Arora, A., Auman, J.T., Ayala, B., et al. (2015). Genomic Classification of Cutaneous Melanoma. Cell *161*, 1681-1696. https://doi.org/10.1016/j.cell.2015.05.044.

10. Miskov-Zivanov, N., Turner, M.S., Kane, L.P., Morel, P.A., and Faeder, J.R. (2013). The duration of T cell stimulation is a critical determinant of cell fate and plasticity. Sci Signal *6*, ra97. 10.1126/scisignal.2004217.

**Table S1. Five examples of interaction attributes used by VIOLIN's decision algorithm.**

| | | Interaction attributes | | Interaction attribute examples | | | | |
|---|---|---|---|---|---|---|---|---|
| | | Attribute name | Attribute symbol | 1 | 2 | 3 | 4 | 5 |
| Elements (nodes, $v$) | Regulator (source node, $v_s$) | Name* | $a_s^{name}$ | CK1 | mTOR | Akt | Resveratrol | RAS |
| | | Type* | $a_s^{type}$ | protein | protein | protein family | chemical | protein family |
| | | Subtype | $a_s^{subtype}$ | kinase | kinase | kinase | antioxidant | GTPase |
| | | HGNC Symbol* | $a_s^{HGNCsymbol}$ | CSNK1A1 | MTOR | AKT1,AKT2, AKT3 | N/A | HRAS, KRAS, NRAS |
| | | Database* | $a_s^{database}$ | UniProt | UniProt | FamPlex | CHEBI | PFAM |
| | | ID* | $a_s^{ID}$ | P48729 | P42345 | AKT | 445154 | PF00071 |
| | | Compartment | $a_s^{compartment}$ | cytoplasm | cytoplasm, nucleus | cytoplasm, membrane | N/A | other |
| | | Compartment ID | $a_s^{compartmentID}$ | GO: 0005737 | GO:0005737, GO:0005634 | GO:0005737, GO:0016020 | N/A | GO:0016020 |
| | Regulated (target node, $v_t$) | Name* | $a_t^{name}$ | APC | Chk1 | GSK3beta | PTEN | p110gamma |
| | | Type* | $a_t^{type}$ | protein | protein | protein | protein | protein |
| | | Subtype | $a_t^{subtype}$ | tumor suppressor | kinase | kinase | phosphatase | enzyme |
| | | HGNC Symbol* | $a_t^{HGNCsymbol}$ | APC | CHEK1 | GSK3B | PTEN | PIK3CG |
| | | Database* | $a_t^{database}$ | UniProt | UniProt | UniProt | UniProt | UniProt |
| | | ID* | $a_t^{ID}$ | P25054 | O14757 | P49841 | P60484 | P48736 |
| | | Compartment | $a_t^{location}$ | cytoplasm | cytoplasm, centrosome | cytoplasm, nucleus, membrane | cytoplasm, nucleus | other |
| | | Compartment ID | $a_t^{locationID}$ | GO: 0005737 | GO: 0005737, GO:0005813 | GO:0005737, GO:0005634, GO:0016020 | GO:0005737, GO:0005634 | GO:0016020 |
| Interaction (edge, $e$) | Influence | Sign* | $a_e^{sign}$ | positive | positive | negative | positive | positive |
| | | Connection Type | $a_e^{connectiontype}$ | direct | indirect | indirect | indirect | indirect |
| | | Mechanism | $a_e^{mechanism}$ | phosphorylation | amount | phosphorylation | transcription | activation |
| | | Site | $a_e^{site}$ | S1504, S150, S1507, S1510 | N/A | S9 | N/A | N/A |
| | Context | Cell Line | $a_e^{cellline}$ | SW480 | cellosaurus:CVCL_0045 | CHO | LNCaP, DU145 | COS-7 |
| | | Cell Type | $a_e^{celltype}$ | colorectal cancer | embryonic | epithelial | prostate cancer | fibroblast-like |
| | | Tissue Type | $a_e^{tissuetype}$ | large intestine | kidney | ovarian | brain, lymph node | kidney |
| | | Organism | $a_e^{organism}$ | human | human | human, mouse, hamster | human | human, monkey |
| | Proven. | Score | $a_e^{score}$ | INDRA: 0.999 | STRING: 0.763, INDRA:0.982 | STRING: 0.996, INDRA:0.999 | STITCH: 0.963, INDRA:0.999 | INDRA:0.997 |
| | | Source | $a_e^{source}$ | literature, expert, database | literature, expert, database | literature, expert, database | literature, expert, database | literature, expert, database |
| | | Statements | $a_e^{statements}$ | Sentence 1 | Sentence 2 | Sentence 3 | Sentence 4 | Sentence 5 |
| | | Paper IDs | $a_e^{paperIDs}$ | PMC2654145, PMID11487578 | PMC4381605 | PMC1403772, PMC3535741 | PMC3181262, PMC2957324 | PMC2652403 |

Sentence examples:
1. "This may be analogous to parallel mechanisms that promote GSK3 phosphorylation of beta-catenin in the absence of Wnt stimulation, such as by GSK3 and CK1 phosphorylation of Axin and APC."
2. "mTOR inhibition in HEK293 cells significantly reduced the total Chk1 level"
3. "Therefore, given that Akt phosphorylates and inactivates GSK3beta, we hypothesized that Akt dependent inactivation of GSK3beta might be responsible for Notch potentiation."
4. "Our results demonstrate that resveratrol induced the expression of PTEN…"
5. "Ras activates p110gamma at the level of the membrane, by allosteric modulation and/or reorientation of the p110gamma…"

Attributes marked with an asterisk (*) are included in the required match conditions.

Sentences from which interaction examples were extracted:

1. "This may be analogous to parallel mechanisms that promote GSK3 phosphorylation of beta-catenin in the absence of Wnt stimulation, such as by GSK3 and CK1 phosphorylation of Axin and APC."
2. "mTOR inhibition in HEK293 cells significantly reduced the total Chk1 level"
3. "Therefore, given that Akt phosphorylates and inactivates GSK3beta, we hypothesized that Akt dependent inactivation of GSK3beta might be responsible for Notch potentiation."
4. "Our results demonstrate that resveratrol induced the expression of PTEN…"
5. "Ras activates p110gamma at the level of the membrane, by allosteric modulation and/or reorientation of the p110gamma…"

**Table S2. Summary of paper retrieval approaches, reader output, interaction list sizes, and VIOLIN runtime for each paper corpus and corresponding interaction list for CS1, CA1.**

| Paper set | Paper retrieval approach | Query | Total papers | Reader | Interaction list | Total interactions | Unique interactions | Unique VIOLIN interactions | Time (s) | Average time (s) |
|---|---|---|---|---|---|---|---|---|---|---|
| $S_{A1}$ | PubMed search | melanoma AND (p70S6K OR S6 OR gsk OR gsk3 OR gsk3a OR gsk3b OR src OR 4ebp1 OR "eIF4E-binding protein 1" OR PHASI OR ybi OR YBX1 OR NSEP1 OR YB1 OR CRD OR Y-box OR SkMel-133) | 6039 | REACH | $R_{A1}$ | 131445 | 105186 | 125545 | 402.254 | 3.06E-03 |
| | | | | INDRA | | 72531 | 49837 | 51628 | 73.3973 | 1.01E-03 |
| | | | | GPT-4.1 | | - | - | - | - | - |
| | | | | Llama 3 | | - | - | - | - | - |
| $S_{A2}$ | REACH Explorer, Fetch | MEK, ERK, AKT, GSK3, P70RSK, S6, CDK4, 4EBP1, YB1, SRC, CHK2, MTOR, PI3K | 120 | REACH | $R_{A2}$ | 6305 | 5200 | 5907 | 3.9387 | 6.25E-04 |
| | | | | INDRA | | 1848 | 1454 | 1524 | 0.8625 | 4.67E-04 |
| | | | | GPT-4.1 | | 2736 | 2640 | 2734 | 1.8008 | 5.91E-04 |
| | | | | Llama 3 | | 5198 | 4920 | 5177 | 3.9539 | 5.65E-04 |
| $S_{A3}$ | REACH Explorer | MAPK/ERK pathway | 50 | REACH | $R_{A3}$ | 1471 | 1352 | 1437 | 0.7782 | 5.29E-04 |
| | | | | INDRA | | 593 | 542 | 559 | 0.3053 | 5.15E-04 |
| | | | | GPT-4.1 | | 1272 | 1255 | 1272 | 0.7994 | 5.33E-04 |
| | | | | Llama 3 | | 1924 | 1853 | 1917 | 1.3654 | 5.86E-04 |
| $S_{A4}$ | REACH Explorer | RPS6K1 | 3 | REACH | $R_{A4}$ | 59 | 51 | 54 | 0.04578 | 7.76E-04 |
| | | | | INDRA | | 30 | 30 | 30 | 0.03521 | 1.17E-03 |
| | | | | GPT-4.1 | | 70 | 70 | 70 | 0.0709 | 1.08E-03 |
| | | | | Llama 3 | | 111 | 111 | 111 | 0.1215 | 2.01E-03 |
| $S_{B1}$ | REACH Explorer | PTEN | 43 | REACH | $R_{B1}$ | 1168 | 944 | 1124 | 0.5304 | 4.54E-04 |
| | | | | INDRA | | 734 | 579 | 619 | 0.3065 | 4.18E-04 |
| | | | | GPT-4.1 | | 816 | 777 | 815 | 0.4603 | 4.74E-04 |
| | | | | Llama 3 | | 986 | 771 | 965 | 0.6159 | 5.57E-04 |
| $S_{B2}$ | Hawse et al. references | N/A | 13 | REACH | $R_{B2}$ | 440 | 403 | 432 | 0.2111 | 4.80E-04 |
| | | | | INDRA | | 380 | 351 | 363 | 0.3065 | 8.07E-04 |
| | | | | GPT-4.1 | | 249 | 245 | 248 | 0.1516 | 4.90E-04 |
| | | | | Llama 3 | | 405 | 382 | 404 | 0.2781 | 7.75E-04 |
| $S_{B3}$ | REACH Explorer | T-cell, PTEN, AKT, FOXO | 11 | REACH | $R_{B3}$ | 373 | 353 | 369 | 0.1834 | 4.92E-04 |
| | | | | INDRA | | 389 | 374 | 384 | 0.181 | 4.65E-04 |
| | | | | GPT-4.1 | | 304 | 299 | 304 | 0.1889 | 5.44E-04 |
| | | | | Llama 3 | | 367 | 332 | 363 | 0.2396 | 7.75E-04 |
| $S_{B*1}$ | REACH Explorer | Breast Cancer, DNA repair, Autophagy, Cancer | 38 | REACH | $R_{B*1}$ | 1223 | 1104 | 1124 | 0.5304 | 4.54E-04 |
| | | | | INDRA | | 847 | 759 | 619 | 0.3065 | 4.18E-04 |
| | | | | GPT-4.1 | | 837 | 814 | 836 | 0.4603 | 4.74E-04 |
| | | | | Llama 3 | | 1111 | 1038 | 1099 | 0.6159 | 5.57E-04 |
| $S_{B*2}$ | REACH Explorer | DNA repair, BRCA1, ADAM17, inflammation | 15 | REACH | $R_{B*2}$ | 871 | 807 | 432 | 0.2111 | 4.80E-04 |
| | | | | INDRA | | 634 | 573 | 363 | 0.3065 | 8.07E-04 |
| | | | | GPT-4.1 | | 274 | 270 | 272 | 0.1516 | 4.90E-04 |
| | | | | Llama 3 | | 483 | 458 | 479 | 0.2781 | 7.75E-04 |

**Total interactions** – interactions retrieved or extracted from database or literature

**Unique interactions** – set of interactions obtained by removing duplicates (interactions with the same: regulator name/id/type, regulated name/id/type, interaction sign) from Total interactions

**Unique VIOLIN interactions** – set of interactions obtained by removing duplicates from Total interactions by VIOLIN (VIOLIN duplicates are interactions that have same elements, context, and influence attributes)

**Time** – time of executing the VIOLIN code with the new interaction list and baseline model files as inputs ($T_T$).

**Average time** – average time per interaction ($^1/_{s_{T,reader}}$)

A.

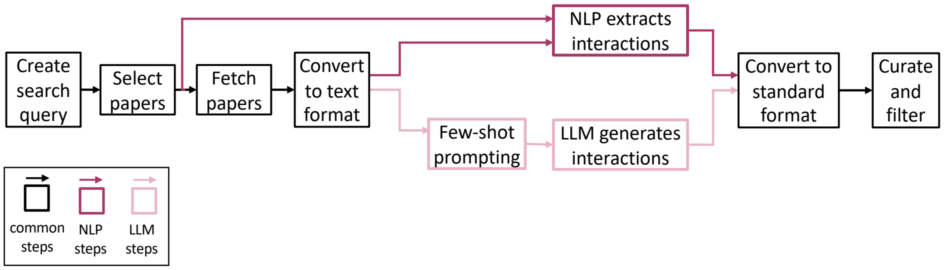


B.

| Legend: -- advantage ✓ – present/required -- drawback x – absent/not req. | | Available features | | | | | Add. steps and resources | | | | | Runtime | |
|---|---|---|---|---|---|---|---|---|---|---|---|---|---|
| Tool | Approach | PubMed access | Paper search | Database access | Open access | API access | Pre-processing | Prompting | Scripts | Parallelization | Large memory | [min/paper] | Device |
| REACH | RB+Stat. | ✓ | ✓ | ✓ | ✓ | ✓ | x | x | x | x | x | ~2 | CPU |
| INDRA | RB+Stat.+ KB | ✓ | ✓ | ✓ | ✓ | ✓ | x | x | x | x | x | ~0.5 | |
| GPT-4.1 | LLM-few shots | x | x | x | x | ✓ | ✓ | ✓ | ✓ | ✓ | ✓ | ~8 | API call |
| Llama 3 | LLM-few shots | x | x | x | ✓ | ✓ | ✓ | ✓ | ✓ | ✓ | ✓ | ~1.9 | GPU |

C.

You are an expert biomedical assistant. Your task is to:
extract relevant information from biomedical research papers and strictly follow the example below,
extract the biological intracellular interaction lists (Biological Regulator, Biological Regulated, Biological Interaction) from the given text.
for Interaction Sign choices: {SIGN_CHOICES},
for Interaction Connection Type choices: {CNX_TYPE_CHOICES},
for Biological Regulator and Biological Regulated Type choices: {TYPE_CHOICES},
for Biological Regulator and Biological Regulated Database choices: {DATABASE_CHOICES},
for Biological Regulator and Biological Regulated Compartment choices: {COMPARTMENT_CHOICES},
for Interaction Mechanism choices: {MECHANISM_CHOICES}.

Q: B-Raf phosphorylates MEK1 and MEK2 on Ser217 and Ser221 , which activates it to dual phosphorylate ERKs , at Thr202 and Tyr204 for human ERK1 and Thr185 and Tyr187 for human ERK2 [ XREF_BIBR , XREF_BIBR ] .

A:
{INTERACTION_1}
{INTERACTION_2}
{INTERACTION_3}
{INTERACTION_4}

Q: Inhibition of NF-kappaB by Gliotoxin , MG132 or Sulfasalazine [ XREF_BIBR ] sensitizes pancreatic cancer cells to apoptosis induced by etoposide ( VP16 ) or doxorubicin .

A:
{INTERACTION_1}
{INTERACTION_2}

Q: {text}

| | |
|---|---|
| SIGN_CHOICES | [positive, negative] |
| CNX_TYPE_CHOICES | [direct, indirect] |
| TYPE_CHOICES | [gene, protein, RNA, chemical, protein family, biological process] |
| DATABASE_CHOICES | [UniProt, HGNC, PubChem, Ensembl, GENCODE ,RefSeq ,Pfam, InterPro, GO, MeSH] |
| COMPARTMENT_CHOICES | [cytoplasm, cytosol, plasma membrane, nucleus, mitochondria ,endoplasmic reticulum ,extracellular] |
| MECHANISM_CHOICES | [phosphorylation, dephosphorylation, ubiquitination, deubiquitination, acetylation, deacetylation, methylation, demethylation, transcription, translation, translocation] |

| | | |
|---|---|---|
| INTERACTION_1 | Biological Regulator:<br>Name: B-Raf<br>Type: protein<br>Subtype: Kinase<br>HGNC Symbol: BRAF<br>Database: uniprot<br>ID: P15056<br>Compartment: N/A<br>Compartment ID: N/A<br><br>Biological Regulated:<br>Name: MEK1<br>Type: protein<br>Subtype: Kinase<br>HGNC Symbol: MAP2K1<br>(Continue to right) | (cont'd)<br>Database: uniport<br>ID: Q02750<br>Compartment: N/A<br>Compartment ID: N/A<br><br>Interaction:<br>Sign: positive<br>Connection Type: direct<br>Mechanism: phosphorylation<br>Site: Ser217/Ser221<br>Cell Line: N/A<br>Cell Type: N/A<br>Tissue Type: N/A<br>Organism: N/A |
| INTERACTION_2 | ... | |
| ... | ... | |

**D.**

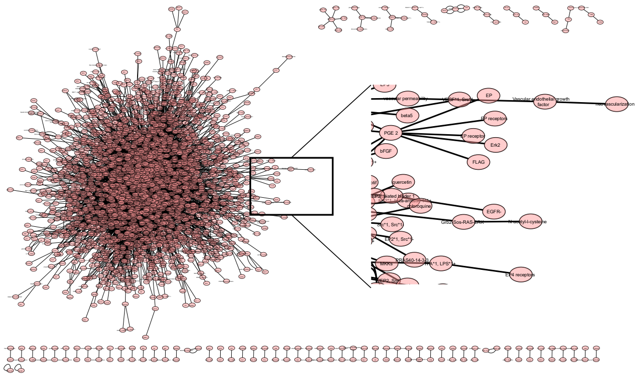


**Figure S3. Overview of paper retrieval methods.**

**A.** The workflow including steps for running rule-based NLP tools and prompting LLMs.

**B**. Summary of the approaches used, advantages and drawbacks of the four reader tools (RB – rule-based, Stat. – statistics, KB – knowledge base). Columns indicate Available features (advantages): programmatic/automated access to PubMed, paper retrieval, and interaction and pathway databases, open-access reader tool, and programmatic (API) access to the tool itself; Additional steps or resources that are required by the tool (disadvantages): paper preprocessing or reformatting before extracting interactions, prompting in order to "train" for suitable output format, additional scripts or code are necessary for running, parallelization for timely output, large memory necessary for processing. Average time per processed paper and the device on which the tool was run.

**C**. Prompt used with ChatGPT-4.1 and Llama3. Prompt comprises a system message specifying the available choices for each attribute and a list of extraction demonstrations, each consisting of a passage paired with a set of annotations. The table on the right presents allowed values for each variable in the prompt.

**D**. Network of interactions in the interaction list $R_{A2}$, which has 2200 nodes and 6305 edges, extracted by REACH for corpus $S_{A2}$.

A. B.

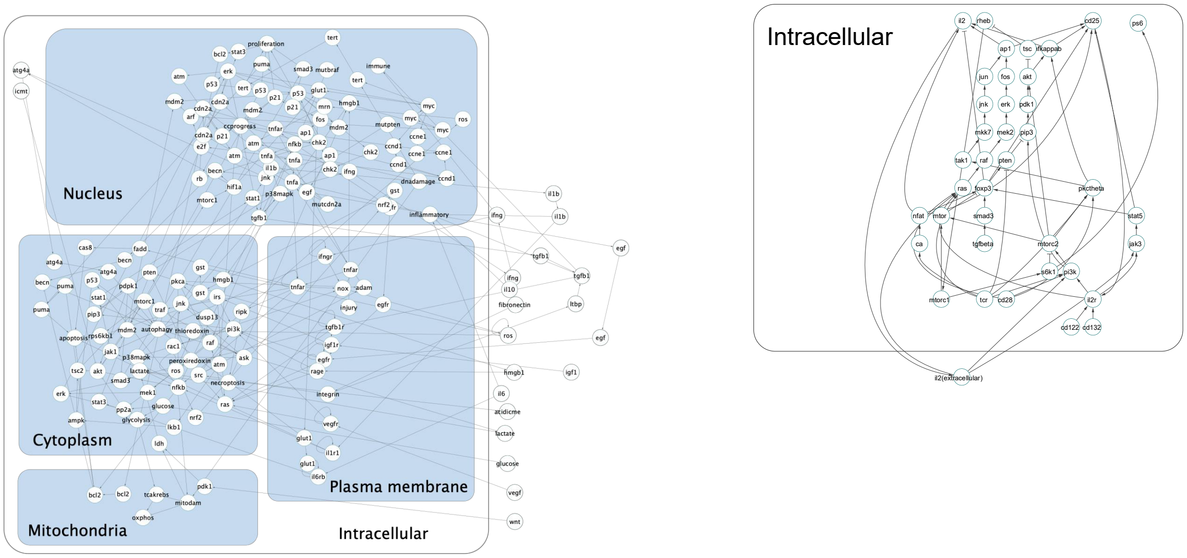


C.

| # | Baseline model | Biological context | Number of nodes | Number of edges | Corresponding corpora |
|---|---|---|---|---|---|
| 1 | **A** | Melanoma cell line SkMel-133 | 179 | 266 | $S_{A1}$, $S_{A2}$, $S_{A3}$, $S_{A4}$ |
| 2 | **B** | Naïve T-cell differentiation to Treg and Th | 39 | 70 | $S_{B1}$, $S_{B2}$, $S_{B3}$, $S_{B*1}$, $S_{B*2}$ |
| 3 | **PID: 8615** | IL23-mediated signaling events | 78 | 38 | $S_{A2}$ |
| 4 | **PID: 11552** | IL27-mediated signaling events | 63 | 26 | $S_{A2}$ |
| 5 | **PID: 22080** | ErbB1 downstream signaling | 287 | 46 | $S_{A2}$ |
| 6 | **PID: 52883** | IL5-medicated signaling events | 56 | 4 | $S_{A2}$ |
| 7 | **PID: 61956** | PDGFR-beta signaling pathway | 374 | 122 | $S_{A2}$ |
| 8 | **PID: 72464** | IFN-gamma pathway | 116 | 29 | $S_{A2}$ |
| 9 | **PID: 73660** | IL6-mediated signaling events | 113 | 39 | $S_{A2}$ |
| 10 | **PID: 81472** | IL12-mediated signaling event | 118 | 36 | $S_{A2}$ |
| 11 | **Reactome: R-HAS-69231** | Cyclin D associated events in G1 | 198 | 25 | $S_{A2}$ |
| 12 | **Reactome: R-HAS-982772** | Growth hormone receptor signaling | 105 | 21 | $S_{A2}$ |
| 13 | **Reactome: R-HAS-1059683** | Interleukin-6-signaling | 70 | 8 | $S_{A2}$ |
| 14 | **Reactome: R-HAS-2559582** | Senescence-Associated Secretory Phenotype (SASP) | 157 | 27 | $S_{A2}$ |
| 14 | **Reactome: R-HAS-2586552** | Signal by Leptin | 52 | 12 | $S_{A2}$ |
| 16 | **Reactome: R-HAS-6785807** | Interleukin-4 and Interleukin-13 signaling | 303 | 29 | $S_{A2}$ |
| 17 | **Reactome: R-HAS-9020956** | Interleukin-27 signaling | 51 | 10 | $S_{A2}$ |

**Figure S4. Networks of baseline graphs A and B.**

**A**. Network of the melanoma cell line SkMel-133, used as baseline graph A.

**B**. Network of the circuitry that controls T cell differentiation we use as a baseline model B with 39 nodes and 70 edges.

**C**. Summary of baseline graphs used, their sizes and corresponding literature corpora. Table lists the IDs and names of graphs that are acquired from external databases, Pathway Interaction Database (PID) and Reactome. Query used to acquire pathway networks through PathwayCommons: “melanoma, BRAF, MEK, ERK, IL6, STAT3, JAK2, CDK4, CDK6”.

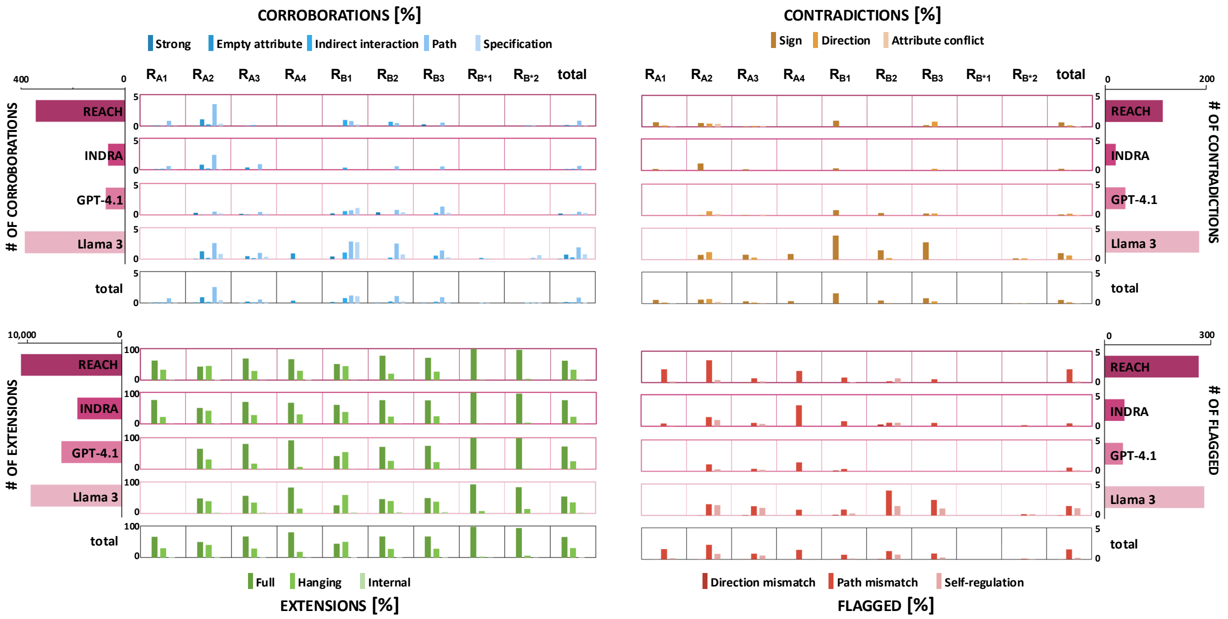


**Figure S5. VIOLIN output for the four reader tools and all nine interaction lists.**

Classification results for all nine interaction lists ($R_{A1-4}$, $R_{B1-3}$, and $R_{B*1-2}$), and all four readers (REACH, INDRA, GPT-4.1 and Llama 3), across all categories (Corroborations, Contradictions, Flagged, and Extensions) and subcategories.

Four category tables (middle): (columns – individual interaction lists) bar heights represent percentages out of total number of interactions in the list for each reader and across all readers; (columns – 'total') bar heights represent percentage for subcategories out of total number of interactions extracted across all lists by an individual reader; (rows – 'total') bar heights represent percentages for subcategories out of total number of interactions for each individual interaction list across all readers; (bottom right corner of each category table) bar heights represent percentages for subcategories out of total number of interactions across all lists and all readers.

Pink bar charts (left and right): the number of interactions extracted by each reader in each category (this excludes list $R_{A1}$ for all readers, since GPT-4.1 and Llama3 were not used on this corpus)

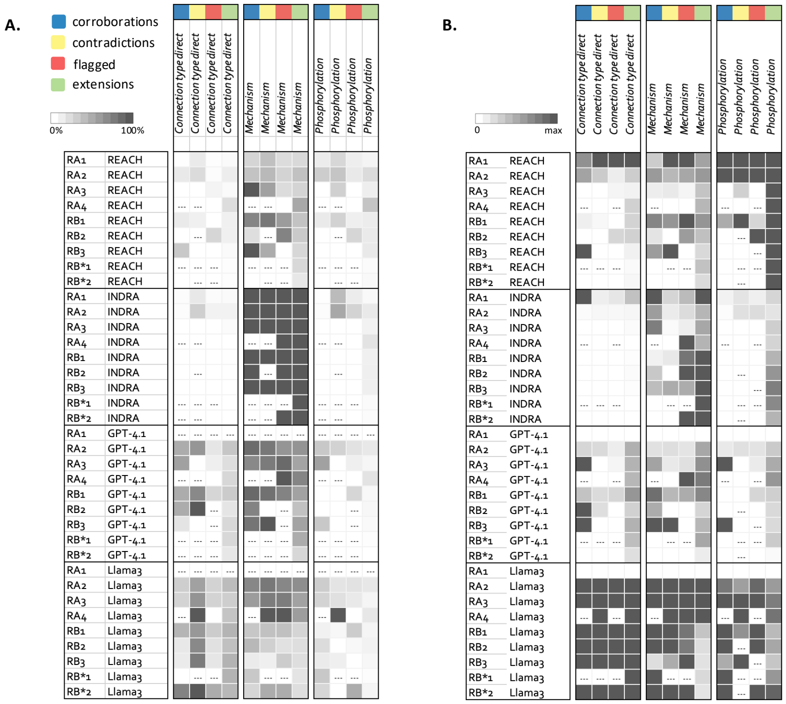

**Figure S6. Attribute coverage for all readers across all corpora and classification categories.**

**A**. Percent presence within extracted interaction list of "direct" in the Connection type attribute, any value for attribute Mechanism, and "phosphorylation" in the Mechanism attribute. Percents are listed relative to each corpus, for each reader.

**B**. Number of interactions with attribute value present for each corpus, each reader, and within each classification category. Maximum number is determined within each category and corpus across all readers. All results are obtained under scheme CS1 and strategy CA4.

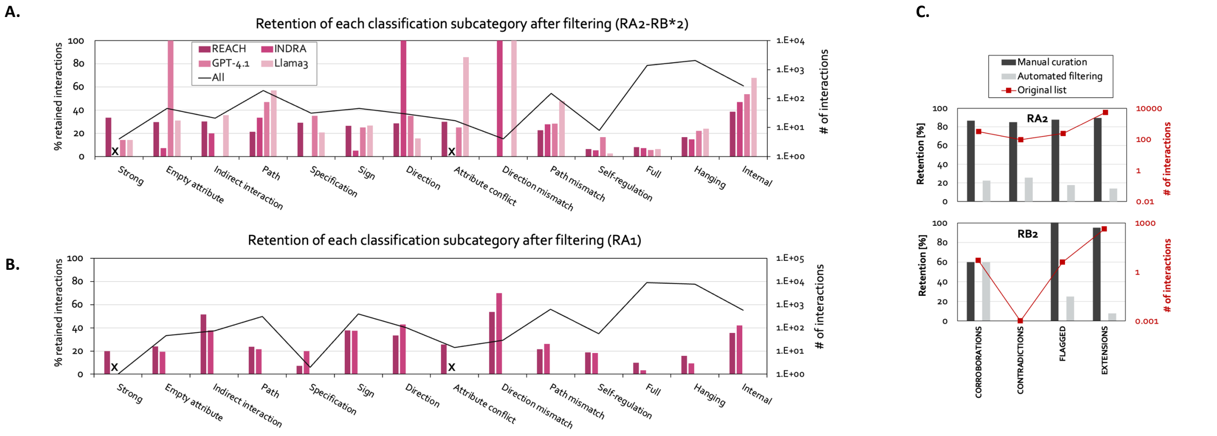


**Figure S7. Retention of interactions after filtering and manual curation.**

**A.** Bars- fraction (%) of interactions retained after filtering shown for each classification subcategory and each reader, computed across corpora $R_{A2}$-$R_{B*2}$. Line- total number of interactions retained after filtering in each subcategory summed across all readers and across corpora $R_{A2}$-$R_{B*2}$.

**B.** Bars- fraction (%) of interactions retained after filtering shown for each classification subcategory for REACH and INDRA and corpus RA1. Line- total number of interactions retained after filtering in each subcategory summed for REACH and INDRA for corpus $R_{A1}$. “x” indicates that there were no interactions in that subcategory before filtering.

**C.** Comparison of FLUTE filtering and manual curation for REACH output, lists $R_{A2}$ and $R_{B2}$.

A.

| Benchmark # | Benchmark Group | Interaction list Description | Interaction list Size | Precision | Recall | Accuracy | F1 |
|---|---|---|---|---|---|---|---|
| 1 | 1 | One from each sub-category | 14 | 1 | 1 | 1 | 1 |
| 2 | 2 | Model A interactions | 266 | 1 | 1 | 1 | 1 |
| 3 | | Model B interactions | 70 | 1 | 1 | 1 | 1 |
| 4 | 3 | A + 10 contradictions | 276 | 1 | 1 | 1 | 1 |
| 5 | | B + 10 contradictions | 80 | 1 | 1 | 1 | 1 |
| 6 | 4 | A + 10 random interactions | 276 | 1 | 1 | 1 | 1 |
| 7 | | B + 10 random interactions | 80 | 1 | 1 | 1 | 1 |
| 8 | 5 | $R_{A2}$-$M_1$ | 2250 | 0.998 | 0.998 | 0.999 | 0.998 |
| 9 | | $R_{A2}$-$M_2$ | 2425 | 0.976 | 0.976 | 0.988 | 0.976 |
| 10 | | $R_{A2}$-$M_3$ | 599 | 0.846 | 0.846 | 0.923 | 0.846 |
| 11 | | $R_{B2}$-$M_1$ | 309 | 1 | 1 | 1 | 1 |
| 12 | | $R_{B2}$-$M_2$ | 94 | 0.989 | 0.989 | 0.9995 | 0.989 |
| 13 | | $R_{B2}$-$M_3$ | 8 | 0.875 | 0.875 | 0.938 | 0.875 |
| 14 | 6 | $R_{A2}$-$M_1$+FLUTE | 222 | 1 | 1 | 1 | 1 |
| 15 | | $R_{A2}$-$M_2$+FLUTE | 405 | 1 | 1 | 1 | 1 |
| 16 | | $R_{A2}$-$M_3$+FLUTE | 171 | 0.977 | 0.977 | 0.988 | 0.977 |
| 17 | | $R_{B2}$-$M_1$+FLUTE | 18 | 1 | 1 | 1 | 1 |
| 18 | | $R_{B2}$-$M_2$+FLUTE | 15 | 1 | 1 | 1 | 1 |
| 19 | | $R_{B2}$-$M_3$+FLUTE | 4 | 1 | 1 | 1 | 1 |

B.

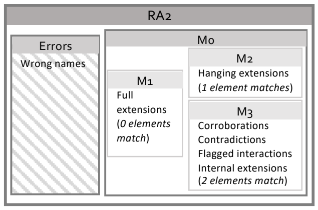


C.

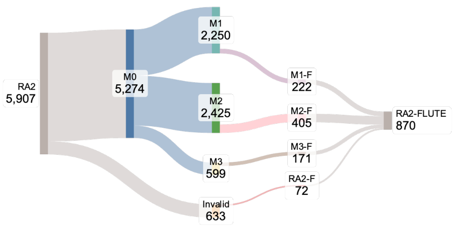


D.

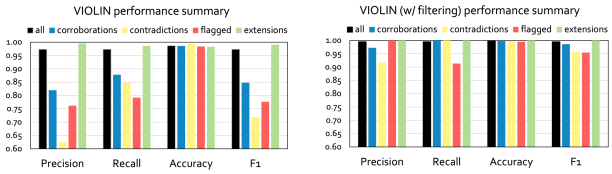


**Figure S8. VIOLIN testing.**

**A.** Summary of tests conducted on VIOLIN. All interactions lists can be found in VIOLIN's GitHub repository.

**B.** Illustration of how tests $M_0$-$M_3$ are organized (shown with RA2, same for RA3).

**C.** Sankey diagram showing interaction list sizes, starting from the original list $R_{A2}$, through list $M_0$, then lists $M_1$, $M_2$, $M_3$, and their corresponding sub-lists after filtering with FLUTE, $M_1$-F, $M_2$-F, $M_3$-F. "invalid" indicates interactions that were removed from $R_{A2}$ with manual curation when creating list $M_0$. The numbers indicate total interaction list size obtained from REACH, before finding redundant interactions.

**D.** Performance summary across all corpora and all readers, for each category, without (left) and with (right) pre-filtering.
